# Towards a neuro-computational account of prism adaptation

**DOI:** 10.1101/187963

**Authors:** Pierre Petitet, Jill X. O’Reilly, Jacinta O’Shea

## Abstract

Prism adaptation has a long history as an experimental paradigm used to investigate the functional and neural processes that underlie sensorimotor control. In the neuropsychology literature, functional explanations of prism adaptation are typically framed within a traditional cognitive psychology ‘box-and-arrow’ framework that distinguishes putative component functions thought to give rise to behaviour (i.e. ‘strategic control’ versus ‘spatial realignment’). However, this kind of theoretical framework lacks precision and explanatory power. Here, we advocate for a computational framework that offers several advantages: 1) an *algorithmic* explanatory account of the computations and operations that drive behaviour; 2) expressed in quantitative mathematical terms; 3) embedded within a principled theoretical framework (Bayesian decision theory, state-space modelling); 4) that offers a means to generate and test quantitative behavioural predictions. This computational framework offers a route toward mechanistic explanations of prism adaptation behaviour. Thus it constitutes a conceptual advance compared to the traditional theoretical framework. In this paper, we illustrate how Bayesian decision theory and state-space models offer principled explanations for a range of behavioural phenomena in the field of prism adaptation (e.g. visual capture, magnitude of visual versus proprioceptive realignment, spontaneous recovery and dynamics of adaptation memory). We argue that this explanatory framework offers to advance understanding of the functional and neural mechanisms that implement prism adaptation behaviour, by enabling quantitative tests of hypotheses that go beyond mere descriptive mapping claims that ‘brain area X is (somehow) *involved* in psychological process Y’.

## 1 Introduction

Adaptation is a fundamental property of the nervous system that enables organisms to flexibly reconfigure sensorimotor processing to counteract perturbations that cause performance errors (Franklin & Wolpert, 2011; Shadmehr, Smith, & Krakauer, 2010). Consider, for example, the case of a basketball player shooting at various times throughout a game. As the game progresses, so muscles will fatigue, such that the same motor command produces a different outcome from one shoot to another. A lateral wind might also alter the trajectory of the ball and deviate it from the aimed basket. In these two situations, an internal (muscle fatigue) or external (wind) disturbance introduces systematic deviations from the intended action goal. These perturbations require the relationship between a desired action goal and the motor commands that execute it to be reconfigured, to avoid the large systematic errors in performance that would ensue if the nervous system were unable to adapt and correct for the perturbations. Thus, adaptation underwrites the maintenance of successful actions across the lifespan.

In a laboratory context, sensorimotor adaptation has been studied experimentally using a variety of methods (e.g. visuomotor rotation, force-field adaptation, saccade adaptation, Coriolis forces, etc.) (Ethier, Zee, & Shadmehr, 2008; Lackner & Dizio, 1994; Mazzoni & Krakauer, 2006; Shadmehr & Mussa-Ivaldi, 1994). Here we focus on a method first developed by von Helmholtz at the end of the nineteenth century, called *prism adaptation* (Von Helmholtz, 1867). In this paradigm, participants wear prism glasses that bend light, and so optically displace the visual field, for example by 10° to the right. When participants perform visuo-motor tasks (e.g. pointing at targets) while wearing the prisms, at first, they make systematic rightward errors (owing to the optical displacement), but participants learn rapidly from the error feedback to correct their movements on subsequent trials and regain normal accuracy (i.e. they adapt). When the prisms are removed post-adaptation, individuals then make errors in the opposite direction, i.e. a leftward “after-effect", which reflects the temporary persistence of some of the compensatory mechanisms engaged during the adaptation. Several features of how prism after-effects generalize or transfer beyond the specifically trained context make it an interesting paradigm to investigate. In healthy controls, prism after-effects tend to generalise at least partially across space (Bedford, 1989, 1993; Gordon M Redding & Wallace, 2006b). This contrasts with visuomotor rotation, for instance, where effects drop off sharply with distance from the trained target location (John W. Krakauer, Pine, Ghilardi, & Ghez, 2000). Pointing during prism exposure is typically aimed at lateral targets under speeded conditions, whereas prism after-effects are often measured at a central (untrained) location, with accuracy emphasized over speed. With this procedure, there is therefore a change in task context between prism exposure and prism after-effect measurement conditions, such that the aftereffect measure intrinsically captures elements of generalization/transfer, at least with respect to task changes (training/test or exposure/after-effect) in reach trajectory, reaching speed and target location. Prism after-effects are measurable in at least three different modalities, visual, proprioceptive, and motor, which appear to have different timescale dynamics (Yohko Hatada, R. Chris Miall, & Yves Rossetti, 2006; Y. Hatada, R.C. Miall, & Y. Rossetti, 2006; Hatada, Rossetti, & Miall, 2006; Gordon M Redding & Wallace, 2001). It has also been claimed that prism aftereffects can transfer to untrained visuospatial tasks (e.g. line bisection task, greyscales task), although these effects in young healthy volunteers appear to occur only with left-shifting (not right-shifting) prisms and to be quite small and variable (Colent, Pisella, Bernieri, Rode, & Rossetti, 2000; Goedert, LeBlanc, Tsai, & Barrett, 2010; Loftus, Vijayakumar, & Nicholls, 2009; Martin-Arevalo, Chica, & Lupianez, 2014; C. Michel, et al., 2003; Schintu, et al., 2014; Striemer, Russell, & Nath, 2016). A stronger evidence base in patients has shown that the after-effects of prism adaptation can transfer to improve cognitive deficits in visuospatial neglect after right hemisphere brain damage (Frassinetti, Angeli, Meneghello, Avanzi, & Làdavas, 2002; Rossetti, et al., 1998; Serino, Barbiani, Rinaldesi, & Làdavas, 2009). After-effects have been shown to transfer to a broad range of untrained sensory and cognitive domains in neglect, including, for example, postural control, occulo-motor exploration, dichotic listening and mental imagery (for review, see: Jacquin-Courtois, et al., 2013). Improved symptomatology after prism adaptation has also been reported in patients with complex regional pain syndrome (Sumitani, et al., 2007) and Parkinson’s disease (Bultitude, Rafal, & Tinker, 2012). This distinctive generalization/transfer profile of prism adaptation, by contrast with other adaptation paradigms, suggests that this experimental model of sensorimotor integration warrants special attention.

What features should a satisfying theoretical account of prism adaptation behaviour have? An ideal account would provide: 1) mechanistic explanations, that are 2) biologically plausible, and 3) can generate quantitative behavioural predictions, 4) about the effect of a range of factors, such as experimental task manipulations (e.g. modality, quality and timing of sensory feedback, gradual versus abrupt perturbation onset, etc.), psychological variables (e.g. internal state estimates of limb position, sensory uncertainty, prior knowledge of the perturbation, etc.), and neural state effects (e.g. change in neural excitability in specific brain region owing to lesion or drug or brain stimulation intervention). Here, we outline the current prevailing (descriptive psychological) model of prism adaptation that is predominant in the literature on healthy individuals, patients and animal studies. We also highlight the brain regions implicated in prism adaptation by studies conceived within this framework. Next, we make the case that a computational characterization of prism adaptation behaviour offers advantages over this traditional functional descriptive approach, and argue the need for an integrated neuro-computational account to further advance understanding within this field.

## 2 Prism adaptation procedures

Since the primary focus of this paper is on prism adaptation experiments, we will first describe how such studies are typically performed. This will provide the reader with the necessary background to engage with the prism adaptation literature and understand the theoretical discussion developed in the following sections.

### 2.1 Typical experimental paradigm

Prism adaptation experiments usually include at least three phases, as follows: 1) <u>pre-adaptation phase</u>: baseline measure of individuals’ visuomotor task performance; 2) <u>prism exposure phase</u>: visuomotor tasks are performed during exposure to the prismatic shift. This shift causes performance errors, and individuals learn to correct their errors gradually and to compensate for the optical shift (i.e. they adapt); 3) <u>post-adaptation phase</u>: baseline tests are repeated, and changes in performance (post - pre) provide a measure of *after-effects* (Kornheiser, 1976) (Figure 1).

**Figure 1.**
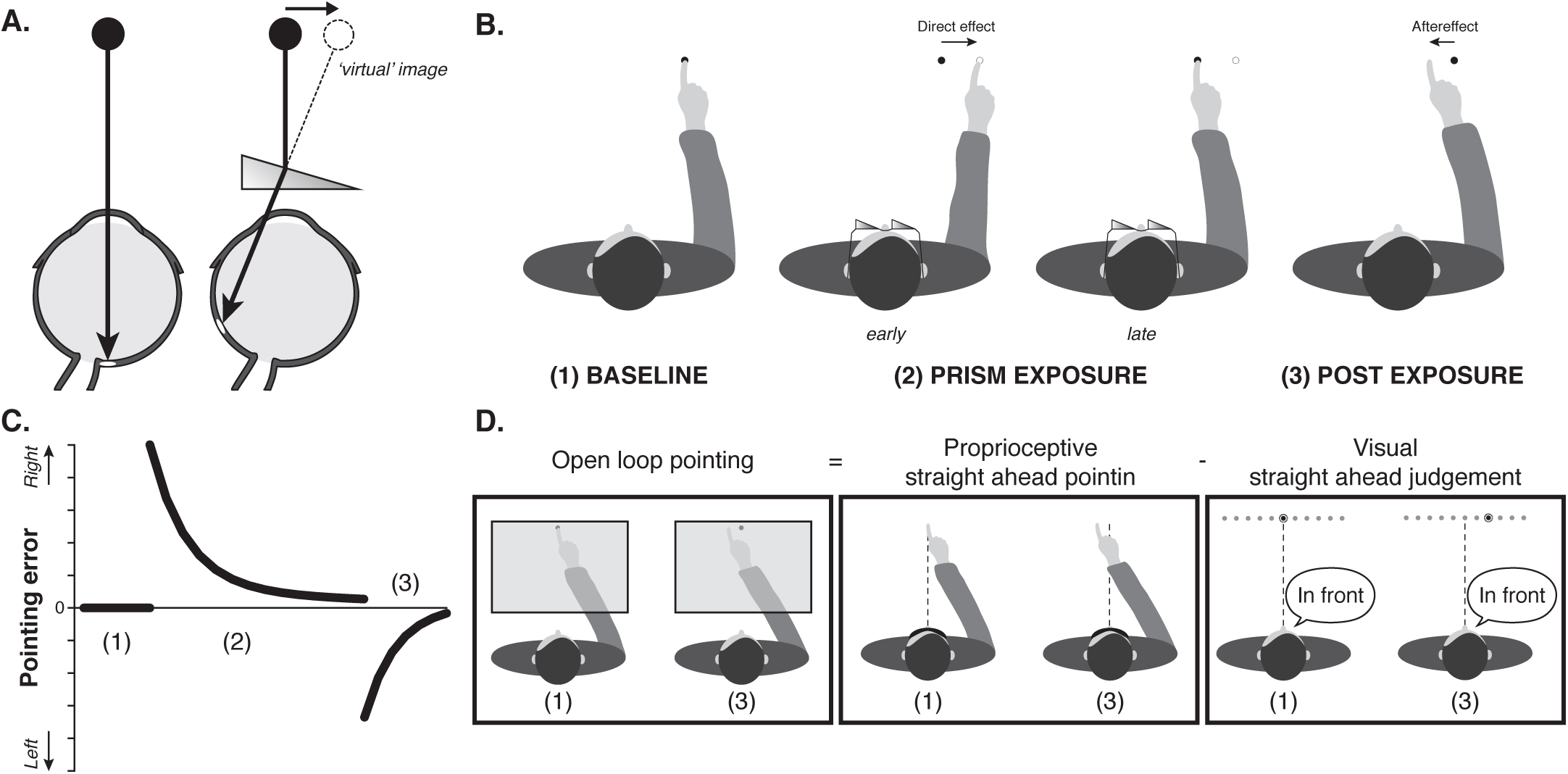
Prism adaptation. **(A)** By bending light, prism lenses displace the visual field in a direction determined by the prism structure. Here for example, light is displaced laterally, by 10° to the right. Hence, a central dot when viewed through this prism is (mis)perceived to be located 10° to the right of its true position. **(B) Typical prism adaptation experimental paradigm**. Participants’ pointing accuracy is first tested at baseline (1), prior to prism exposure. Figure illustrates closed-loop pointing at baseline, i.e. participant is required to make fast and accurate pointing movements to a visual target and receives visual feedback of the reach trajectory and endpoint. During prism exposure (2), the goggles shown in A) are worn. Owing to the optical shift, the ‘direct effect’ is that the participant makes rightward pointing errors initially (early phase), but learns gradually from trial-by-trial error feedback to correct these errors and re-gain baseline pointing accuracy (late phase). Consequent leftward prism after-effects (errors) are measurable post-adaptation once the glasses have been removed (3). **(C) Canonical pattern of performance errors during prism adaptation**. Plot shows reach endpoint error (y-axis) as a function of trial number (x-axis) during closed loop pointing (i.e.: with visual feedback). Note baseline accuracy (i.e. mean error centred on zero) (1), followed by rightward errors (in the direction of the prismatic shift) that decrease gradually across prism exposure trials (2), followed by leftward errors (in the direction opposite the prismatic shift) after removal of the prism goggles (3). **(D) Three tests commonly used in the prism adaptation literature to quantify prism after-effects**. During *open loop pointing* participants point at visual targets, which are viewed transiently, and visual feedback of the reach trajectory and the reach endpoint is deprived. This prevents (further) learning from endpoint error (which would over-turn the after-effect). *Open-loop pointing* measures of after-effect are deviated in the direction opposite the prismatic shift (i.e. here leftward). During *proprioceptive straight ahead pointing* blindfolded participants are asked to point in the direction they perceive as being straight ahead of their nose. This is thought to capture the proprioceptive component of adaptation. This after-effect measure is also deviated in the direction opposite to the prismatic shift (i.e. leftward). During *visual straight ahead judgement* participants must indicate when a flashing light is perceived as being straight ahead of their nose. This is thought to capture the visual component of adaptation. After-effects with this measure are deviated in the same direction as the prismatic shift (i.e. rightward). The sum of visual and prioprioceptive after-effects immediately after prism exposure has been shown to equal the magnitude of after-effect quantified by open-loop pointing, which is therefore known as the *total visumotor shift* (Y Hatada, et al., 2006; Hay & Pick Jr, 1966; Gordon M Redding & Wallace, 1988; Gordon M. Redding & Wallace, 1996; Templeton, et al., 1974).

In the studies we will consider, participants are typically asked to make pointing movements during each of the three phases. Visual feedback of individuals’ trial-by-trial reach endpoints can either be provided (closed loop pointing, CLP) or deprived (open loop pointing, OLP). During prism exposure (phase 2), endpoint errors are initially deviated in the direction of the optical displacement on both of these types of pointing. Performance change relative to baseline (during - pre) during this initial phase is often described as the *direct effect* of prisms. If visual feedback of endpoint errors is provided during prism exposure (i.e. closed loop pointing), individuals tend to reduce these errors progressively to regain baseline accuracy (i.e. they adapt) (Figure 1B, C).

Prism after-effects are measured by asking participants to point again after removal of the prisms. If sufficient practice has occurred during prism exposure, performance will now be deviated in the direction opposite to the prismatic shift. If visual feedback of these (now leftward) reach endpoints is provided (i.e. closed loop pointing), performance errors will once again be corrected rapidly (now in the opposite direction) to re-gain baseline accuracy (Figure 1C). This de-adaptation (or washout) can be limited by depriving visual feedback (i.e. using open loop pointing) (Figure 1D). Prism after-effects measured using open loop pointing can persist even after closed loop performance has returned to baseline levels (Inoue, et al., 2015).

Two other tests are commonly contrasted across the baseline and post-exposure test phases, to measure modality-specific prism after-effects: visual straight-ahead judgements and proprioceptive straight-ahead pointing (Y. Hatada, R. Miall, & Y. Rossetti, 2006; Yohko Hatada, Yves Rossetti, et al., 2006; Gordon M Redding & Wallace, 1988; Gordon M. Redding & Wallace, 1993; Gordon M Redding & Wallace, 2001). The visual straight ahead judgement requires participants to report verbally when a visual stimulus moving laterally across their visual field is perceived as being straight ahead of their nose (Figure 1D). This measure is thought to rely on eye-head coordination, and any post-exposure change in accuracy is usually interpreted as a visual after-effect (Y Hatada, et al., 2006). Proprioceptive straight ahead pointing requires blindfolded participants to point to the position in space they perceive to be straight ahead of their nose. It is thought to provide a measure of head-hand coordination, and post-exposure shifts in this measure are interpreted as a proprioceptive after-effect (C. S. Harris, 1963; Templeton, Howard, & Wilkinson, 1974). Several authors have reported that immediately after prism adaptation, the sum of the visual and proprioceptive after-effects – measured by visual straight ahead judgement and proprioceptive straight ahead pointing, respectively – equals the magnitude of after-effects assayed using open loop pointing (Y Hatada, et al., 2006; Hay & Pick Jr, 1966; Gordon M Redding & Wallace, 1988; Gordon M. Redding & Wallace, 1996; Templeton, et al., 1974). This finding is relatively intuitive given that both vision of target location at onset and proprioceptive and motor feedback during the reach trajectory contribute to motor performance on open loop pointing. Open loop pointing quantifies the combined contribution of both factors influencing hand-eye coordination (i.e. the total visuomotor shift), while visual straight ahead judgement and proprioceptive straight ahead pointing each measure individual components (eye-head and head-hand respectively). Studies vary in whether they study just the total visuomotor shift, or the visual or proprioceptive after-effects, or investigate the relationship between all three.

### 2.2 Important factors to consider

Several experimental factors can strongly influence the modality, magnitude and persistence of both the direct effects and the after-effects of prism adaptation. The way the prismatic shift is introduced (gradually versus abruptly), the visibility of the starting position of the limb, the availability of visual feedback during the movement trajectory versus only at the reach endpoints (i.e. concurrent versus terminal exposure), the duration of prism exposure, the movement speed, the target location, or the limb used - all have been shown to influence behavioural performance (Bedford, 1989; Hamilton, 1964; Inoue, et al., 2015; Kitazawa, Kimura, & Uka, 1997; Carine Michel, Pisella, Prablanc, Rode, & Rossetti, 2007; Gordon M. Redding & Wallace, 1996; Gordon M Redding & Wallace, 2006a). Additionally, brain lesions can affect the way individuals adapt to prisms and express after-effects (Bossom, 1965; Weiner, Hallett, & Funkenstein, 1983; Welch & Goldstein, 1972).

In the following section, we will outline the descriptive theoretical framework typically used within the neuropsychology literature to explain the effects of the various factors listed above. Subsequently, we will argue for the advantages of a computational framework in place of this descriptive account. Key benefits of this formal model framework are that it offers: 1) principled mechanistic explanations of behaviour, 2) which specify the computations that give rise to behaviour, 3) in precise mathematical terms, 4) that enable quantitative tests of behavioural predictions, 5) and characterize information processing in terms (mathematical functions) that could plausibly be implemented by neural circuits (unlike the traditional box-and-arrow cognitive psychology model). We argue that this explanatory framework offers a significant conceptual advance, which promises to accelerate progress in understanding the causal bases of prism adaptation behaviour, in terms of the algorithms that drive it, the neural circuits that implement it, and how these interact.

## 3 The traditional dual-process framework: strategic control versus spatial realignment

### 3.1 Theoretical framework

For the past forty years, the large majority of studies investigating the neural mechanisms underlying prism adaptation have interpreted their results within a theoretical framework that distinguishes two learning processes that contribute differentially to the direct effects (i.e.: error correction) versus the after-effects of prisms. This framework posits that prism adaptation recruits two distinct functional mechanisms: a rapid process of error reduction that reflects strategic adjustments in motor control, and a slower process, so-called ‘true’ sensorimotor adaptation, thought to reflect the spatial realignment of motor, proprioceptive and visual coordinate reference frames (for review, see: Gordon M Redding, Rossetti, & Wallace, 2005; Gordon M Redding & Wallace, 2006b). The name given to these two processes has varied slightly across studies and over time, but the core idea of a distinction between a fast strategic component and a slower ‘true’ sensorimotor realignment has remained consistent. Here we will adopt Redding and Wallace’s (Gordon M Redding, et al., 2005; Gordon M Redding & Wallace, 2006b) most recent terminology to describe this dual-process theoretical framework and review the evidence for functional and anatomical dissociations of these two processes.

Within this framework, the constituent processes driving prism adaptation are described as follows. *Strategic control* is a set of processes that guide everyday adaptive motor behaviours. For example, to reach for a cup, depending on the sensory information available, it is necessary to first select the appropriate reference frame to code the target location (e.g. visual-motor, proprioceptive-motor) and guide the appropriate reach-to-grasp command. The process of setting the reference frame in relation to the desired action is called ‘calibration’ (Gordon M Redding, et al., 2005; G. M. Redding & Wallace, 2002). Strategic control also requires selecting the region of extrapersonal space most relevant to the on-going task (e.g. the shelf where the cup is located), and is therefore described as being closely related to spatial attention. If a reach-to-grasp action is unsuccessful, the action may have to be ‘recalibrated’ by various means. For example, (while wearing prisms) one may direct his/her reaching movement to the side of the cup so as to reduce the previous motor error. This mix of partly automatic, partly conscious processes is thought to contribute predominantly to the rapid error correction that occurs during the initial phase of prism exposure, but to contribute poorly to the prism after-effects (Aimola, Rogers, Kerkhoff, Smith, & Schenk, 2012; L. Pisella, Michel, Grea, Tilikete, & Rossetti, 2004; Gordon M Redding, et al., 2005; Gordon M Redding & Wallace, 2006b; Weiner, et al., 1983).

It is important to note that, within this framework, strategic motor control adjustments do not require the linkage between different coordinate frames to be reconfigured. Instead, motor performance can be improved simply by changing the motor command issued to reach the target. An analogy commonly used to illustrate this idea is to consider a rifleman with misaligned telescopic sight. Suppose that the scope of the rifle is misaligned with the barrel, so that the marksman misses the target systematically by 10° to the right. Simply re-aiming 10° to the left of the target would allow the marksman to hit his target without having to realign the scope with the barrel. Strategic control refers to this process of quickly recalibrating the reference frame to meet the task objectives. Successful movements driven by such a process should therefore be specific to the trained task context and should not generalize to other contexts (e.g. other movements, other spatial locations, etc.).

By contrast, *spatial realignment* refers to the process of adjusting for constant differences in spatial coordinates between multiple sensorimotor coordinate systems. In reference to the previous analogy, this would be equivalent to realigning the barrel with the scope of the rifle. The traditional theoretical framework often posits that accurate pointing movements require the correct alignment of two reference frames: an eye-head visual-motor system (measured with the visual straight ahead test) and a head-hand proprioceptive-motor system (measured with straight ahead pointing) (Gordon M Redding, et al., 2005; G. M. Redding & Wallace, 2002). Because the prismatic shift displaces the visual-motor reference frame only, compensatory shifts in the visual-motor and/or proprioceptive-motor systems are required in order to re-align the two reference frames and regain correct eye-hand coordination during prism exposure (Templeton, et al., 1974). The temporary carry-over of these adaptive shifts after removal of prisms is thought to be predominantly responsible for the after-effects of prism adaptation.

### 3.2 Behavioural predictions

In summary, the core theses of the traditional dual-process framework are that: 1) strategic recalibration is a cognitively demanding process, that drives error correction early during prism exposure, but contributes little to prism after-effects; 2) spatial realignment is an automatic process, that develops more gradually, and is mainly responsible for prism aftereffects; 3) these two processes operate relatively independently from one another (Figure 2). Within this theoretical framework, closed loop performance during prism exposure reflects both the strategic control and spatial realignment components, whereas prism after-effects (measured via open loop pointing, visual straight ahead judgement or proprioceptive straight ahead pointing) provide a measure of spatial realignment. Impaired strategic control would therefore affect error correction during prism exposure, but would not directly affect prism after-effects. By contrast, impaired spatial realignment would affect both direct effects and after-effects of prisms.

**Figure 2.**
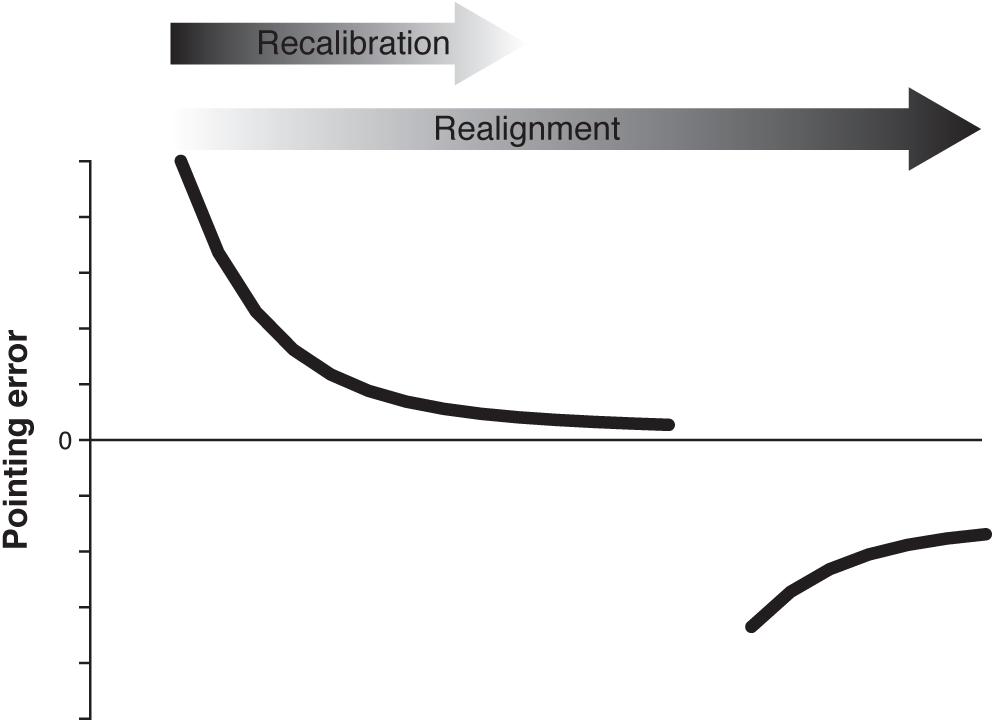
The traditional dual-process framework. The traditional theoretical framework posits that error correction during prism adaptation relies on two processes. *Strategic control* refers to the calibration of individuals’ task workspace around the task relevant objects. It is described as a rapid process that intervenes early during prism exposure but contributes poorly to prism after-effects. Conversely, *spatial realignment* is described as developing more slowly during prism exposure and is thought to be responsible for the prism after-effects. The term describes a process of bringing the different sensorimotor coordinate frames (visual-motor, proprioceptive-motor) into alignment with each other.

In the following two sections (3.3 and 3.4), we will review some of the main empirical evidence that supports this theoretical model (for in depth review, see: Gordon M Redding, et al., 2005; G. M. Redding & Wallace, 2002; Gordon M Redding & Wallace, 2006b) and summarise attempts to localize the neural correlates of the two proposed processes.

### 3.3 Strategic control

Within the traditional framework, strategic control refers to the process of quickly recalibrating the reference frame to (re)code the location of the target. It is thought to contribute mainly to the early phase of prism exposure.

Consistent with this view, several studies have reported behavioural markers of a learning component engaged during the early phase of prism exposure, but saturating quickly, and contributing poorly to the after-effects. For example, in healthy individuals, during the initial phase of prism exposure, when errors are large, the time gap between target foveation and onset of the pointing movement increases transiently, but returns to normal within about 10 trials, as the endpoint error magnitude is gradually reduced (Rossetti, Koga, & Mano, 1993). A similarly rapid timecourse was observed in trial-by-trial corrections of the initial acceleration phase of the reach trajectory during the first 10 prism exposure trials (O’Shea, Gaveau, Kandel, Koga, & Susami, 2014). These behavioural data are consistent with a rapid error corrective process engaged early during prism exposure. It has also been shown that imposing a cognitive load (mental arithmetic) during prism exposure disrupts participants’ ability to correct pointing errors while wearing prisms (Gordon M Redding, Rader, & Lucas, 1992). Taken together, these results implicate a strategic learning component that contributes to error correction during early prism exposure. None of these behavioural phenomena have been shown to relate quantitatively to measures of prism after-effects.

The neural substrates associated with this ‘strategic control’ component of prism adaptation have been inferred from brain lesions that lead to impaired error reduction during prism exposure, with spared after-effects (Canavan, et al., 1990; Fernandez-Ruiz, et al., 2007; Newport & Jackson, 2006; L. Pisella, et al., 2004; Weiner, et al., 1983; Welch & Goldstein, 1972) (Table 1). Such studies have converged on crucial involvement of the cerebral cortex in strategic control (Canavan, et al., 1990; Weiner, et al., 1983; Welch & Goldstein, 1972), more specifically, the posterior parietal cortex (Newport & Jackson, 2006; L. Pisella, et al., 2004). The functional specificity implied by this behavioural pattern is debatable given the variety of neurological impairments that can lead to this pattern of performance. For example, temporal lesions (Canavan, et al., 1990), spinocerebellar ataxia type 2 (Fernandez-Ruiz, et al., 2007), and spatial neglect (Aimola, et al., 2012) have all been associated with impaired error reduction but normal prism after-effects.

**Table 1.**
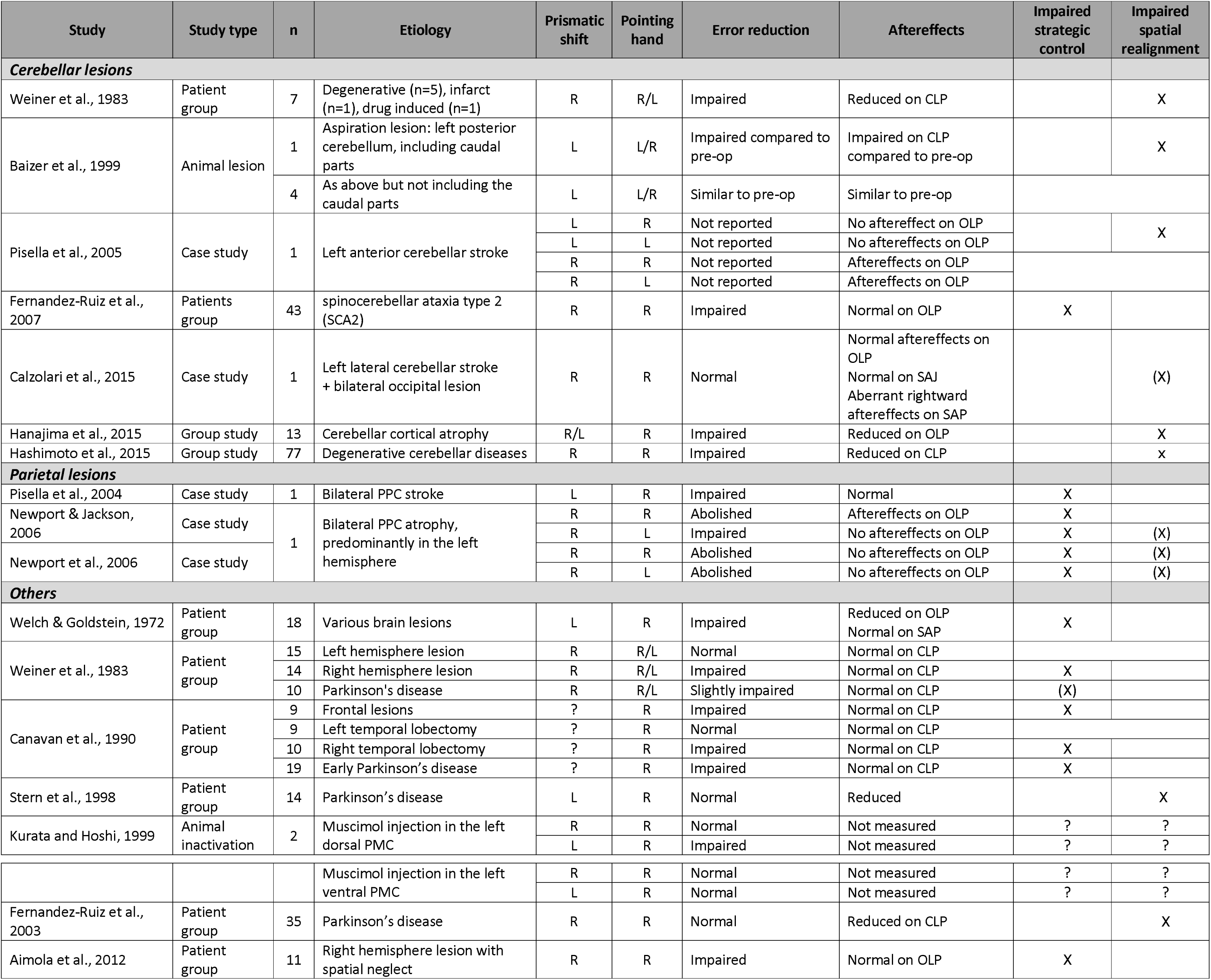
Overview of the lesion studies of prism adaptation. This table includes the main human and animal lesion studies of prism adaptation and is not meant to be exhaustive. PMC = Pre-motor Cortex; PPC = Posterior Parietal Cortex; CLP = Closed Loop Pointing; OLP = Open Loop Pointing; SAP = Straight Ahead Pointing; SAJ = Straight Ahead Judgement

| Study | Study type | n | Etiology | Prismatic shift | Pointing hand | Error reduction | Aftereffects | Impaired strategic control | Impaired spatial realignment |
| --- | --- | --- | --- | --- | --- | --- | --- | --- | --- |
| **Cerebellar lesions** |  |  |  |  |  |  |  |  |  |
| Weiner et al., 1983 | Patient group | 7 | Degenerative (n=5), infarct (n=1), drug induced (n=1) | R | R/L | Impaired | Reduced on CLP |  | X |
| Baizer et al., 1999 | Animal lesion | 1 | Aspiration lesion: left posterior cerebellum, including caudal parts | L | L/R | Impaired compared to pre-op | Impaired on CLP compared to pre-op |  | X |
|  |  | 4 | As above but not including the caudal parts | L | L/R | Similar to pre-op | Similar to pre-op |  |  |
| Pisella et al., 2005 | Case study | 1 | Left anterior cerebellar stroke | L | R | Not reported | No aftereffect on OLP |  | X |
|  |  |  |  | L | L | Not reported | No aftereffects on OLP |  |  |
|  |  |  |  | R | R | Not reported | Aftereffects on OLP |  |  |
|  |  |  |  | R | L | Not reported | Aftereffects on OLP |  |  |
| Fernandez-Ruiz et al., 2007 | Patients group | 43 | spinocerebellar ataxia type 2 (SCA2) | R | R | Impaired | Normal on OLP | X |  |
| Calzolari et al., 2015 | Case study | 1 | Left lateral cerebellar stroke + bilateral occipital lesion | R | R | Normal | Normal aftereffects on OLP<br>Normal on SAJ<br>Aberrant rightward aftereffects on SAP |  | (X) |
| Hanajima et al., 2015 | Group study | 13 | Cerebellar cortical atrophy | R/L | R | Impaired | Reduced on OLP |  | X |
| Hashimoto et al., 2015 | Group study | 77 | Degenerative cerebellar diseases | R | R | Impaired | Reduced on CLP |  | x |
| **Parietal lesions** |  |  |  |  |  |  |  |  |  |
| Pisella et al., 2004 | Case study | 1 | Bilateral PPC stroke | L | R | Impaired | Normal | X |  |
| Newport & Jackson, 2006 | Case study | 1 | Bilateral PPC atrophy, predominantly in the left hemisphere | R | R | Abolished | Aftereffects on OLP | X |  |
|  |  |  |  | R | L | Impaired | No aftereffects on OLP | X | (X) |
| Newport et al., 2006 | Case study |  |  | R | R | Abolished | No aftereffects on OLP | X | (X) |
|  |  |  |  | R | L | Abolished | No aftereffects on OLP | X | (X) |
| **Others** |  |  |  |  |  |  |  |  |  |
| Welch & Goldstein, 1972 | Patient group | 18 | Various brain lesions | L | R | Impaired | Reduced on OLP<br>Normal on SAP | X |  |
| Weiner et al., 1983 | Patient group | 15 | Left hemisphere lesion | R | R/L | Normal | Normal on CLP |  |  |
|  |  | 14 | Right hemisphere lesion | R | R/L | Impaired | Normal on CLP | X |  |
|  |  | 10 | Parkinson's disease | R | R/L | Slightly impaired | Normal on CLP | (X) |  |
| Canavan et al., 1990 | Patient group | 9 | Frontal lesions | ? | R | Impaired | Normal on CLP | X |  |
|  |  | 9 | Left temporal lobectomy | ? | R | Normal | Normal on CLP |  |  |
|  |  | 10 | Right temporal lobectomy | ? | R | Impaired | Normal on CLP | X |  |
|  |  | 19 | Early Parkinson's disease | ? | R | Impaired | Normal on CLP | X |  |
| Stern et al., 1998 | Patient group | 14 | Parkinson's disease | L | R | Normal | Reduced |  | X |
| Kurata and Hoshi, 1999 | Animal inactivation | 2 | Muscimol injection in the left dorsal PMC | R | R | Normal | Not measured | ? | ? |
|  |  |  |  | L | R | Impaired | Not measured | ? | ? |
|  |  |  | Muscimol injection in the left ventral PMC | R | R | Normal | Not measured | ? | ? |
|  |  |  |  | L | R | Normal | Not measured | ? | ? |
| Fernandez-Ruiz et al., 2003 | Patient group | 35 | Parkinson's disease | R | R | Normal | Reduced on CLP |  | X |
| Aimola et al., 2012 | Patient group | 11 | Right hemisphere lesion with spatial neglect | R | R | Impaired | Normal on OLP | X |  |

In the healthy brain, several groups have used neuroimaging techniques to try and identify brain regions involved in strategic control during prism adaptation. Typically, these studies have contrasted the amplitude of neural activity in the ‘early’ versus ‘late’ phases of prism exposure and interpreted areas activated by this contrast (i.e. early > late) as neural correlates of the strategic control component of prism adaptation (Table 2). Consistent with lesion studies, activity in the posterior parietal cortex has been reliably reported by whole brain neuroimaging studies, but other brain regions such as the primary motor cortex, the anterior cingulate cortex, and the cerebellum have also been found to be preferentially activated during the early versus late phase of prism exposure (Clower, et al., 1996; Danckert, Ferber, & Goodale, 2008; Kuper, et al., 2014; Luauté, et al., 2009) (Table 2). Aside from functional localization in this way, these neuroimaging studies have said little about *how* these identified brain regions are thought to implement the process of strategic control.

**Table 2.**
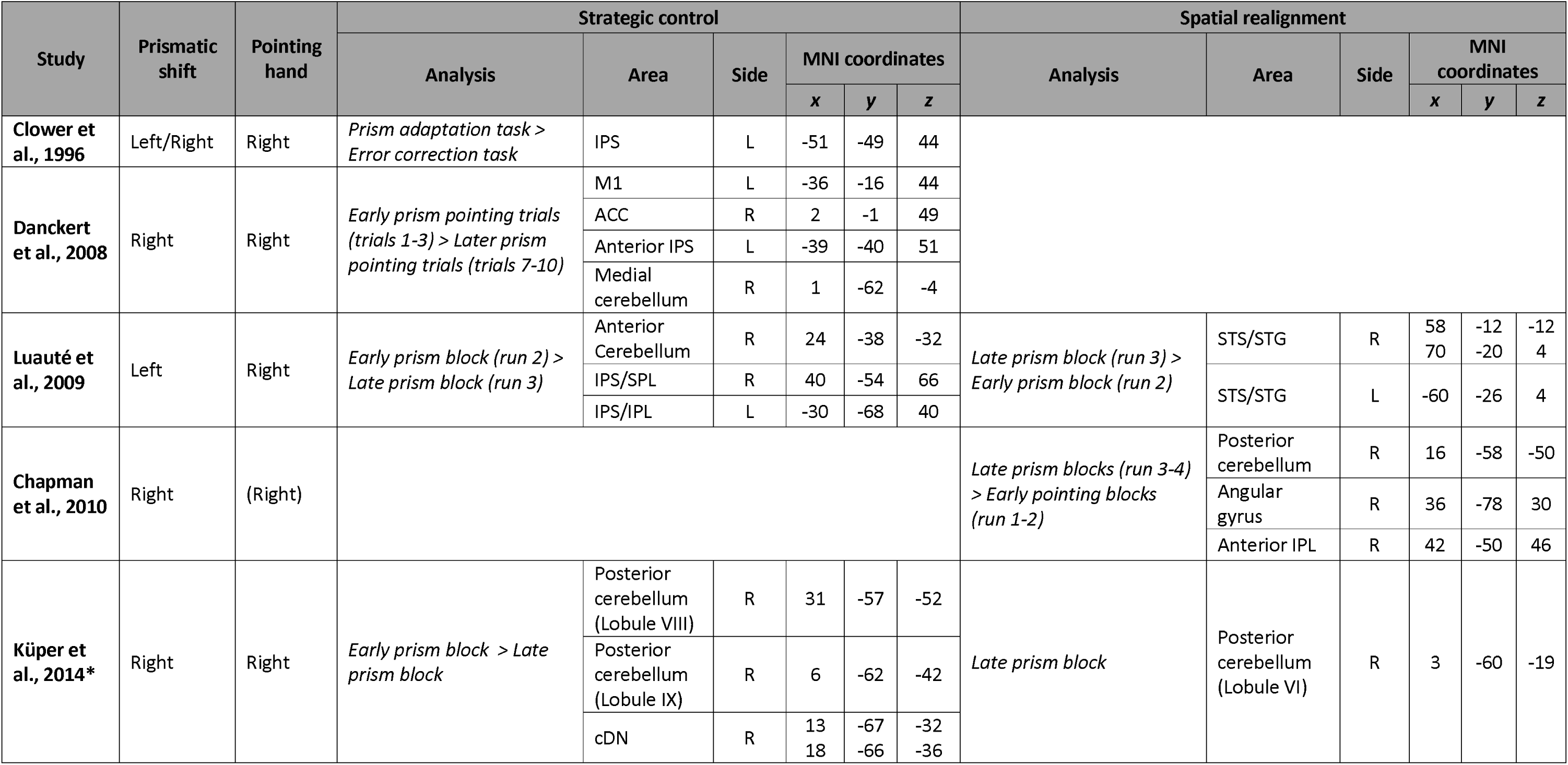
Overview of the neuroimaging studies of prism adaptation. * Study including the cerebellum and dentate nucleus only. IPS = Inferior Parietal Sulcus; M1 = Primary Motor Cortex; ACC = Anterior Cingulate Cortex; SPL = Superior Parietal Lobule; cDN = caudate Dentate Nucleus; STS/STG = Superior Temporal Sulcus/ Superior Temporal Gyrus

### 3.4 Spatial realignment

Within the traditional dual process theoretical framework, spatial realignment refers to the process of shifting the visual-motor and/or proprioceptive-motor systems in order to realign these two coordinate frames and regain accurate hand-eye coordination. This process is believed to develop slowly during prism exposure, and to be responsible almost entirely for the prism after-effects (Gordon M Redding, et al., 2005; G. M. Redding & Wallace, 2002; Gordon M Redding & Wallace, 2006b).

One study in healthy individuals reported a putative kinematic signature of sensorimotor realignment: gradual correction of the terminal (deceleration) phase of the pointing trajectory during prism exposure. This corrective process unfolded slowly during prism exposure and the magnitude of correction of this kinematic error correlated with the magnitude of prism after-effects (O’Shea, et al., 2014). Endpoint error appears not to be necessary for spatial realignment to occur, as after-effects can be observed in the absence of measurable reach endpoint errors if the prismatic shift is introduced gradually (Dewar, 1971; Hanajima, et al., 2015; Howard & Freedman, 1968; Carine Michel, et al., 2007). Instead, it has been suggested that the discordance between the expected (feedforward predicted) and observed (feedback measured) position of the hand is the learning signal for spatial realignment (Gordon M Redding & Wallace, 2006b). Support for this claim comes mainly from the finding that the magnitude of visual and proprioceptive after-effects (measured by the visual straight ahead judgement and proprioceptive straight ahead pointing, respectively) depends upon the sensory information available in-flight during the reach trajectory when individuals are wearing prisms (Gordon M. Redding & Wallace, 1996; Gordon M Redding & Wallace, 2001). If both proprioceptive and visual feedback is available, proprioceptive aftereffects tend to be greater than visual after-effects. The opposite is true (i.e. greater visual than proprioceptive after-effects) if only proprioceptive feedback is available (Gordon M Redding & Wallace, 2001). This suggests that the modality in which the discrepancy between the predicted and observed hand location is sensed determines which coordinate reference frame is preferentially re-aligned.

Several studies have reported evidence of impaired spatial realignment following cerebellar lesions in humans (Calzolari, Bolognini, Casati, Marzoli, & Vallar, 2015; Hanajima, et al., 2015; Martin, Keating, Goodkin, Bastian, & Thach, 1996; L Pisella, et al., 2005; Weiner, et al., 1983) (Table 1) and non-human primates (Baizer, Kralj-Hans, & Glickstein, 1999). The typical behavioural pattern is reduced error correction during prism exposure, combined with decreased or absent prism after-effects. The contribution of the precise cerebellar sub-regions is still unclear, as evidence of impaired spatial realignment has been found after anterior (Calzolari, et al., 2015; L Pisella, et al., 2005) and posterior (Baizer, et al., 1999; Martin, et al., 1996) lesions to cerebellum. To our knowledge, only three neuroimaging studies have investigated the pattern of functional brain activity associated with spatial realignment (Chapman, et al., 2010; Kuper, et al., 2014; Luauté, et al., 2009). They did so by investigating brain regions that were more active during the later stage of prism exposure compared to the early stage (late > early). Posterior cerebellar activity was reported in two of these studies (Chapman, et al., 2010; Kuper, et al., 2014), but other regions such as the superior temporal gyrus and angular gyrus were also activated (Chapman, et al., 2010; Luauté, et al., 2009) (Table 2).

### 3.5 Is this theoretical framework satisfying?

The traditional dual-process theoretical framework offers an account of various behavioural dissociations observed in healthy individuals and neurological patients (for review, see: Gordon M Redding, et al., 2005; G. M. Redding & Wallace, 2002; Gordon M Redding & Wallace, 2006b). The main functional insight provided by this framework has been to distinguish two psychological processes (strategic control, spatial realignment) that combine to explain behaviour, but seem to operate with a certain degree of independence from one another. However, studies conceived within this theoretical framework do not offer a mechanistic explanation of how prism adaptation behaviour arises, and attempts to localise the neural circuits underlying these two processes have yielded heterogeneous results (Table 1). In addition, the information processing operations executed by the identified neural components remain largely unknown.

The traditional dual-process theory suffers from a pervasive problem in cognitive neuroscience – how to bridge the conceptual gap between cognitive psychological level descriptions of behaviour and biologically plausible descriptions of neural circuit functioning? Computational models offer a potential bridge, as they provide a common currency (algorithms, mathematical functions) in which to describe both information processing and neural circuit mechanisms that could implement such functions. We contend that, in order to advance cognitive neuroscience explanation of the causal brain-behaviour dynamics that underwrite prism adaptation, the traditional dual-process framework needs to be replaced with a re-conceptualization at the algorithmic level. By ‘algorithmic’, we mean a level of description that sets out clearly the (mathematical) rules and operations (functions) required to execute prism adaptation behaviour. It is obvious that, at the cellular and neural circuit level, cognitive concepts like ‘strategic control’ or ‘spatial realignment’ have no explanatory value, since brain circuits are computing information, and the explanatory task is to provide an account of how these computations implement psychological functions. Explanatory progress therefore requires a conceptual advance: a theoretical framework that decomposes prism adaptation behaviour into the underlying algorithms required to implement it (for a related recent argument, see: John W Krakauer, Ghazanfar, Gomez-Marin, MacIver, & Poeppel, 2017). Re-conceptualizing prism adaptation in terms of its constituent algorithms offers objective mathematical description, as opposed to qualitative description offered by the traditional dual-process psychological framework. This greater precision helps avoid confusion related to terminology. It also allows for quantitative experimental predictions. We will return to these points in the next section. More generally, recasting any behaviour or cognitive process in algorithmic terms helps advance the field towards a re-defined taxonomy of cognitive processes, one grounded in the recognition that the same computations (and brain circuits) might contribute to diverse behavioural phenomena depending on the constraints of the task. This offers a way to move beyond accounts of brain activations in terms of ‘area X is involved in cognitive function Y (e.g. attention)’ to ‘computation Y is *implemented* within circuit X and engaged during tasks a, b, c’. This distinct conceptual approach could, for example, help to explain why prism after-effects transfer from pointing tasks to cognitive domains in neglect patients (Rossetti, et al., 1998; Sumitani, et al., 2007).

In the following section, we propose an algorithmic decomposition of the computations required for prism adaptation, by leveraging insights from the field of computational neuroscience applied to sensorimotor control. We argue that this decomposition, while not incompatible with the traditional psychological approach, offers a more useful theoretical framework, in terms of quantitative precision, explanation and plausible neural implementation.

## 4 Computational principles of sensorimotor control

### 4.1 The temporal dynamics of adaptation explained by multiple timescale models

State-space modelling, has provided insights into the temporal dynamics contributing to motor behaviour during sensorimotor adaptation (Smith, Ghazizadeh, & Shadmehr, 2006). Trial-by-trial error correction has been explained as the output of multiple adaptive systems that learn and forget on different timescales. The key idea is that these systems compete to learn from performance error and that the sum of their states provides an inverse estimate of the perturbation that is used to correct motor performance (Figure 3). A commonly used simplified model posits that the multitude of possible timescales can be approximated by a *fast system*, which learns and forgets rapidly, and a *slow system*, which learns and forgets more slowly (Smith, et al., 2006). The model predicts that early during adaptation, error correction is mainly dominated by the fast system, which saturates quickly before decaying back to baseline. By contrast, the slow system develops more gradually over extended practice and accounts for most of the error correction that occurs during later stages of exposure to the perturbation. This ‘two-state’ framework is able to reproduce and explain counterintuitive behavioural phenomena in adaptation, such as spontaneous recovery, that is the re-appearance of aftereffects after a brief period of washout (Lee & Schweighofer, 2009; Smith, et al., 2006; Zarahn, Weston, Liang, Mazzoni, & Krakauer, 2008). It offers an efficient way to extract (hidden) temporal dynamics of learning processes driving adaptation and to generate precise quantitative predictions based on the relative contributions of the fast versus slow systems to motor behaviour during adaptation. For example, it has been shown that the level of adaptation reached by the slow system during force-field adaptation, rather than the overall performance improvement, predicts the amount of long-term retention (Joiner & Smith, 2008).

**Figure 3.**
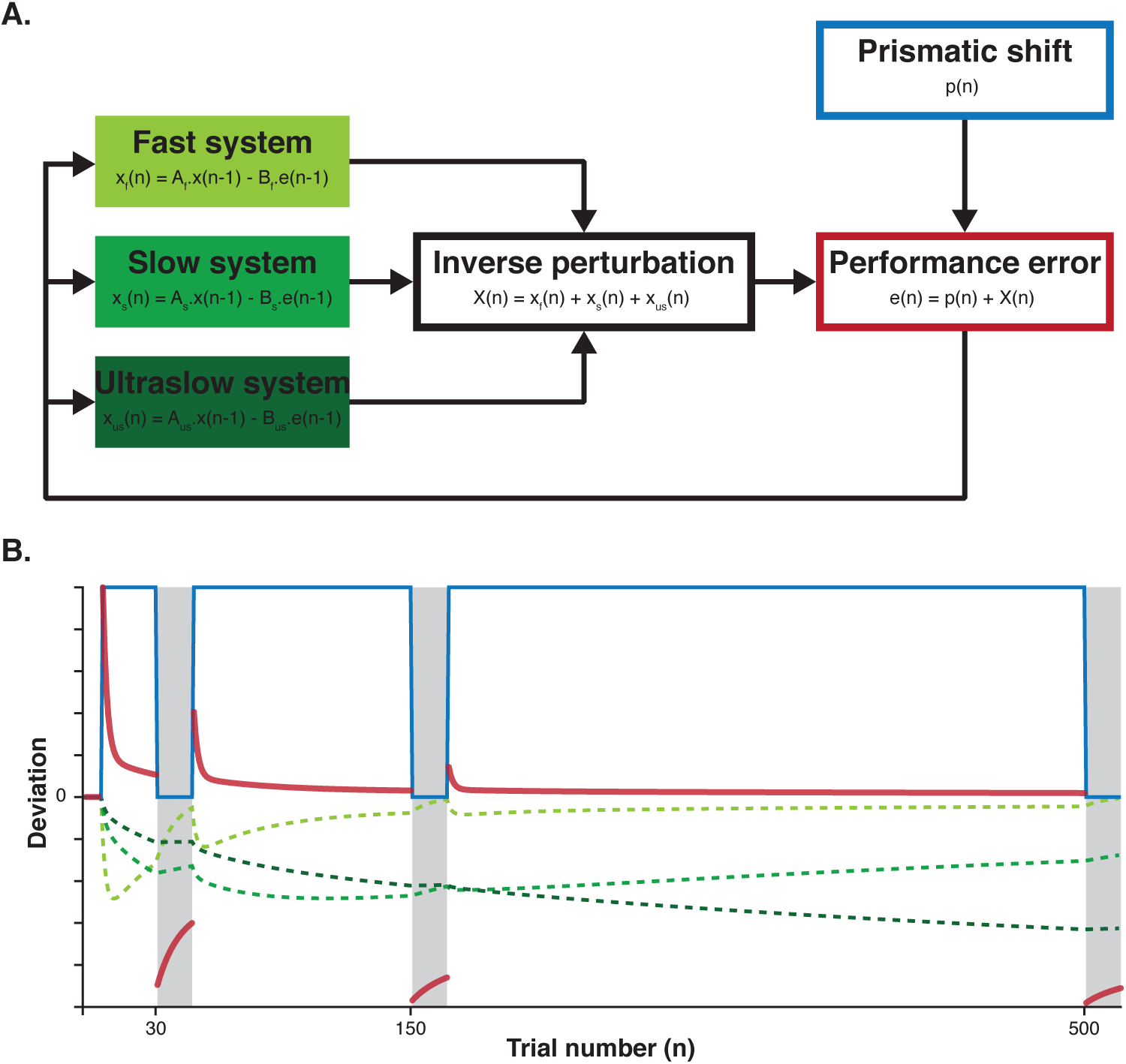
The three timescales state-space model for prism adaptation. **(A)** Each of the fast, slow and ultraslow systems are described by a pair of free parameters: a retention factor A, that describes the amount of decay occurring between trials (0 <A _f_ < A_s_ < A_us_ < 1), and a learning rate B, that describes the fraction of performance error being incorporated into that system’s state on each trial (0 < B_us_ < B_s_ < B_f_ < 1). The sum of the states of the three systems produces an inverse estimate of the perturbation on a trial-by-trial basis, which can be used to induce a change in motor output. On any trial, performance error therefore corresponds to the sum of the prismatic shift and the state of the three systems. We have chosen here to illustrate state-space models with a three timescales model because of its relevance for prism adaptation (Inoue, et al., 2015) but in principle any number of systems could be posited. **(B)** Simulation of prism adaptation in the three timescales model, based on Inoue and colleagues’ experiment (2015). When the prismatic shift (in blue) is introduced, the three systems (dotted green lines) learn at three different rates (set by their respective B). The sum of the states of the three systems is added to the magnitude of the prismatic shift to reduce performance error (in red) during the exposure. If the after-effects are probed with open loop pointing after 30 exposure trials (learning rate is set to 0 for all three systems on open loop pointing), the memory trace decays rapidly because it is largely dominated by the fast system, which has the lowest retention factor. However, if prism exposure continues, the contribution of the fast system gradually decreases. As a result, after-effects are more stable after 150 trials. Finally, after extended exposure of 500 trials, the ultraslow system accounts for most of the error correction. Because of that system’s high retention factor, the after-effects are then very stable.

Because they are not restricted to two learning systems (Inoue, et al., 2015; Kim, Ogawa, Lv, Schweighofer, & Imamizu, 2015; K. P. Kording, Tenenbaum, & Shadmehr, 2007), multiple timescales models can also be used to ask what number of processes combine to best explain adaptation behaviour. For example, a recent prism adaptation study asked how many systems were needed to explain the immediate and long-term retention of prism after-effects (on open loop pointing). The authors found that two systems were sufficient to explain their data for brief duration prism exposure (< 150 pointing trials) but that a third ‘ultraslow’ system was required when exposure was more prolonged (500 trials). This result is in accordance with findings from the saccadic adaptation literature showing that the brain’s estimate of the likely timescale over which a perturbation is occurring (temporal credit assignment) determines the timescale over which the memory is subsequently retained (K. P. Kording, et al., 2007). In other words, there may be a continuum of learning processes, operating over multiple timescales that vary with factors such as the source, duration, and volatility of the perturbation. Following similar reasoning, a recent study combined state-space modelling and functional magnetic resonance imaging to test for neural signatures associated with a range of possible learning timescales (30 were posited) contributing to adaptation to a novel visuomotor rotation (Kim, et al., 2015). In this study, a dimensionality reduction analysis revealed four main components (i.e. group of multiple timescales), each associated with different neural networks and learning dynamics. Notably, all these networks included either a parietal and/or a cerebellar component, suggesting that the activation timecourses of brain regions known to be associated with sensorimotor adaptation are compatible with the state-space modelling framework. In a different field, recent advances in biophysical models of neural networks also support the idea that different brain networks process information at different timescales (Bernacchia, Seo, Lee, & Wang, 2011; Chaudhuri, Knoblauch, Gariel, Kennedy, & Wang, 2015; Fusi, Drew, & Abbott, 2005). This may offer a potential path to bridge biologically plausible models of neural networks with computational descriptions of learning systems and span explanatory levels from neurons to behaviour.

In summary, computational models including multiple learning timescales offer a good quantitative description of motor behaviour during sensorimotor adaptation. The main theoretical limitation of this approach is its agnosticism regarding the information content that is being learnt (and forgotten) by these multiple systems during adaptation. Prism adaptation induces visuo-proprioceptive conflict that requires the nervous system to modify sensory (visual, proprioceptive) and motor information processing to regain normal behavioural accuracy. What is the relationship between these functional components (visual, proprioceptive and motor systems) and the learning dynamics extracted by state-space models? The following sections outline a computational framework (Bayesian decision theory) that clarifies the content of information processing in relation to the visual, proprioceptive and motor systems.

### 4.2 Internal models for sensorimotor control

Prior to generating a successful action (e.g. pointing to a visual target), the nervous system needs to produce a coherent, accurate and unbiased estimate of the current state of one’s body, the external world, and how they interact, based on all useful sources of information available. Based on this knowledge, an action plan must then be selected, one that is most likely to maximise performance in relation to the current behavioural goal. The selection of the most appropriate action plan is fundamentally a decision process that is tightly coupled to the ability to generate accurate predictions about the consequences of one’s actions on the world and/or one’s body (Konrad P Körding & Wolpert, 2006).

Within this framework, sensorimotor control is proposed to rely on internal models of the external world and the mechanics of the body. Inverse models transform a desired behavioural goal into an action plan to accomplish it (Kawato, 1999; Daniel M Wolpert & Kawato, 1998). When applied to the sensory domain, inverse models infer the current state of the body (e.g. position of the hand in space) from sensory input (e.g. proprioceptive feedback). Forward models work in the opposite direction and predict the next state (e.g. next position of the hand) based on an estimate of the current state, a copy of the motor command (efference copy), and some internal representation of the complex causal relationship between the two (for review, see: Davidson & Wolpert, 2005; Franklin & Wolpert, 2011; Lalazar & Vaadia, 2008; Miall & Wolpert, 1996; Shadmehr, et al., 2010). In the context of this paper, we will distinguish between a visual and a proprioceptive forward model, each of which generates predictions about the likely next (i.e.: expected) visual and proprioceptive state. One advantage of continuously predicting the next state of the body is that this limits the impact of neural transmission delays inherent in relying on actual sensory feedback instead of feedforward predictions (Shadmehr, et al., 2010). It also allows the brain to compare continuously the veridicality of its predictions against the actually observed (i.e. sensed) measure of a state (provided by sensory inverse models). Deviations between predicted and observed states (i.e.: prediction errors) can be used as a signal to drive updating of internal models (see section 4.3.3).

### 4.3 Bayesian decision theory

In this section, we outline a theoretical framework that offers a mathematical description of the concepts introduced above (in section 4.2). Bayesian decision theory is composed of two components, Bayesian statistics and decision theory. Bayesian statistics offers a means to formalise how an ideal observer should combine new information (e.g. observed sensory input) with prior beliefs (e.g. predicted sensory input), and how multiple sources of uncertain information (e.g. multiple sensory modalities, predictions, prior knowledge) should be integrated, in an *optimal* fashion, to generate a more certain combined estimate of the current state. Decision theory describes the process of rationally selecting actions, based on the predictions of internal models, given the current behavioural goal. Bayesian decision theory therefore provides a general framework to formalize optimal state estimation and action selection in a dynamic and uncertain world.

#### 4.3.1 Decision theory and rational movement planning

Any given action (e.g. grasping a cup, hitting a ball) can be executed by an almost infinite number of possible movements (Figure 4). Yet, individuals often move in a very stereotypical way (C. M. Harris & Wolpert, 1998; Morasso, 1981; Simmons & Demiris, 2005). Why is this so? Decision theory provides a mathematical framework that formally explains why individuals choose to move the way they do. The central concept is that the cost of a potential movement (e.g. energy consumed, fatigue, risk of an injury, etc.) is weighed against the potential reward that is expected from that movement (e.g. sporting success, monetary gain, altruistic feelings, etc.) (Mazzoni, Hristova, & Krakauer, 2007; Shadmehr, Huang, & Ahmed, 2016). Mathematically, the concept of a *utility function* (or *cost function*, the negative counterpart) captures the complex relationship that integrates all of these factors, and quantifies the overall desirability of a potential movement choice. Within this decision theoretic framework, the process of selecting one action amongst alternatives is operationalized as the rational choice of whichever movement plan maximizes expected utility (or minimizes expected cost) (M. Berniker & Kording, 2011; Konrad P Körding & Wolpert, 2006). Mathematically, expected utility is defined as:

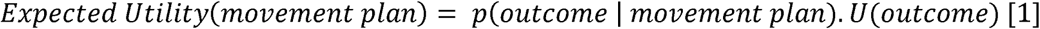

where *p(outcome | movement plan)* is the current estimate of the probability of obtaining an outcome given a particular movement plan (e.g. probability of reaching the target given a certain aiming direction) and *U(outcome)* is the utility associated with this outcome. This mathematical definition becomes intuitive if you consider, for example, a gambling task in which individuals have to choose between alternative options that are associated with varying reward probabilities and reward magnitudes (for example: Behrens, Woolrich, Walton, & Rushworth, 2007; Hsu, Bhatt, Adolphs, Tranel, & Camerer, 2005). In this scenario, the expected utility of each action alternative is the probability of that action yielding a reward, multiplied by the reward magnitude. Choosing the option that maximizes expected utility is the definition of choosing rationally. This mathematical framework has been shown to provide a good quantitative fit to human behavioural data in this kind of decision making task (Behrens, et al., 2007; Hsu, et al., 2005) and has been used to identify functional brain imaging signals that co-vary with the computation of these decision variables (Daw, O’doherty, Dayan, Seymour, & Dolan, 2006; Hampton, Bossaerts, & O’doherty, 2006; John P. O’Doherty, Dayan, Friston, Critchley, & Dolan, 2003).

**Figure 4.**
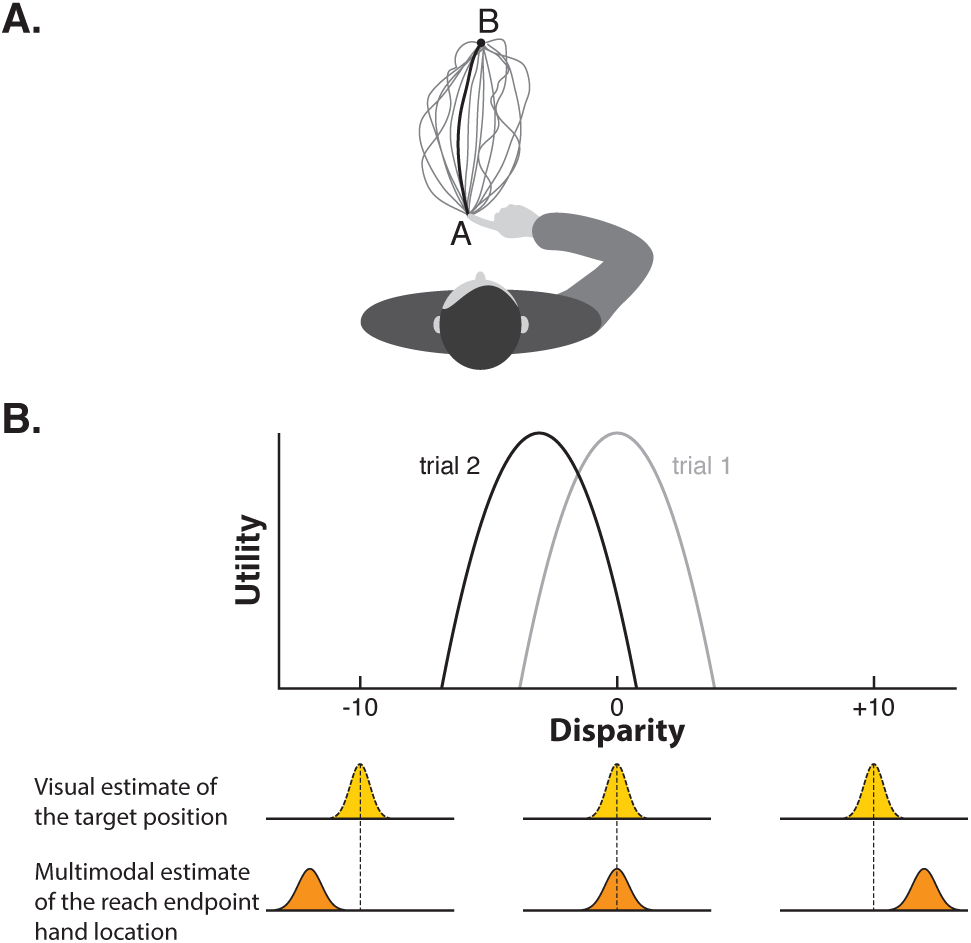
Decision theory. **(A)** Any action can be executed in an almost infinite number of ways. For example, many different movement trajectories would bring one’s finger from point A to a point B. Decision theory provides a mathematical framework that describes how a rational decision maker should choose among alternative movement parameters based on their relative level of expected utility. **(B)** Prism adaptation can be conceptualized as a manipulation that affects the computation of expected utility associated with the disparity between the visual estimate of the position of the target and the multimodal estimate of the hand location at the reach endpoint. Before prism onset, the movement plan with highest expected utility is the one that minimises this disparity (i.e. peak utility centred on zero on trial 1). After prism onset however, the visuo-proprioceptive conflict introduced by the prisms induces a performance error: the (experienced) utility of the executed movement plan doesn’t match the predicted utility. This should result in a shift of utility in the direction opposite to the error when planning the pointing movement on trial 2: now, the movement plan with highest expected utility is one that results in a negative disparity between the visual estimate of the target position and the multimodal estimate of the hand location at the reach endpoint (i.e. the multimodal estimate of the hand location is to the left of the visual estimate of the target).

When planning a movement, more than one decision needs to be made (Figure 4). For example, reaching to point at a visual target requires selection of an aiming location (under prism displacement this need not overlap with the estimated target position see Figure 4B), as well as movement speed and movement trajectory. These different aspects of movement planning are likely to have different utility functions, and decision theory offers a quantitative framework within which these can be integrated to determine rational movement decisions. Whereas in the gambling task example, the experimenter imposes a utility function, by setting the reward structure of the task, in real-life motor control contexts, it is hypothesised that an agent’s rational choice of movements is guided by internal utility functions. Assuming that individuals naturally choose actions that maximize expected utility, then investigating the rules that dictate individuals’ movement properties offers a window onto the internal utility functions that guide action choices. When generating target directed movements, for example, it has been shown that healthy individuals favour the precision rather than smoothness of movement (C. M. Harris & Wolpert, 1998), so as to adopt a trajectory that minimizes what resembles the terminal squared error (Konrad Paul Körding & Wolpert, 2004). Movement speed seems to be determined by both a speed-accuracy trade-off and an implicit cost assigned to the metabolic energy consumed to produce the movement (Mazzoni, et al., 2007; Shadmehr, et al., 2016).

Within the decision theory framework, factors that change expected utility should result in a change of movement plan. The prism adaptation task can be conceptualized as a manipulation that changes the expected utility of pointing movements. Based on equation 1, there are two ways of modifying the expected utility: 1) by changing the utility associated with certain movement outcomes (*U(outcome))*, and/or 2) by changing the probability of obtaining a certain outcome given a movement plan (*p(outcome | movement plan)*). These two could be argued to map on to the traditional dual-process model distinction between strategic control and spatial realignment, respectively.

Strategic control can be re-conceptualized as changing *U(outcome)*, where the outcome is defined as the disparity between the estimate of the hand position at the reach endpoint and the estimate of the target position. Normally, peak utility is when this disparity is zero, i.e. the pointing finger lands on the same location as the (visual) target. However, when an individual corrects strategically for a rightward displacement, by choosing not to aim at the perceived (right-shifted) target location, but instead to aim left of that perceived location, s/he is effectively defining a new utility function with its peak at a non-zero disparity (Figure 4B). Evidence that movement plans are sensitive to changes in *U(outcome)* has been found in an experiment that imposed new utility functions by modulating the monetary value associated with certain movement outcomes (Trommershäuser, Maloney, & Landy, 2003). Under these imposed task constraints, healthy individuals were able to rationally select movement endpoints that maximized the potential reward.

*Spatial realignment* can be conceptualized, within this framework, as minimization of the prediction error associated with the expected *p(outcome | movement plan)*. That is, the prism manipulation changes the probability that a planned pointing movement (with a given aiming location) will accurately hit the target (*p(hitting the target | aiming location)*). Because of the optical shift, during the early phase of prism exposure the experienced utility of pointing movements does not match predictions (*U(missing the target by 10° to the right*) < *U(hitting the target)*), i.e. there is a prediction error in terms of utility. We propose that this signal should induce changes in internal models in order to update the computation of *p(outcome aiming location)*. This would gradually displace the peak of expected utility towards a different movement plan (i.e. more left-oriented) to achieve the desired (unchanged) outcome (i.e. point accurately at the target).

It is worth noticing that this explanatory framework can thus incorporate the key feature of the traditional theoretical framework, i.e. the dissociation between two alternative ways to reduce motor errors during prism exposure (Gordon M Redding, et al., 2005; Gordon M Redding & Wallace, 2006b) (see section 3.1).

A potential criticism of this proposed decision theory framework is that it merely re-describes the traditional dual-process model and does not add anything new to the understanding of prism adaptation. The answer to this objection is threefold. First, there is value in precise quantitative (mathematical) description of algorithms that explain behaviour, by contrast with qualitative description. Re-specifying behaviour at an algorithmic level also offers a means to generate and test hypotheses about the neural implementation of those computations (John W Krakauer, et al., 2017; John P O’doherty, Hampton, & Kim, 2007). Second, the model we propose offers a formal mathematical description of how ‘strategic control’ and ‘spatial realignment’ processes interact, which is typically vague or lacking in the traditional framework. Third, this framework incorporates the contribution of both task errors (i.e. breaches of expectancy in terms of the utility of a movement) and reinforcement learning (Sutton & Barto, 1998) in driving behavioural change during adaptation (Huberdeau, Krakauer, & Haith, 2015; John W Krakauer, et al., 2017). A recent adaptation experiment using visuomotor rotation demonstrated this interaction, by showing that rates of adaptation and retention were differentially modulated by monetary rewards and punishments associated with movement outcomes (Galea, Mallia, Rothwell, & Diedrichsen, 2015). Thus changes in expected utility differentially affected the rate of error correction and the magnitude of subsequent after-effects. Since patients with Parkinson’s disease are likely to have altered utility functions caused by a depletion in dopamine in the basal ganglia (Frank, Seeberger, & O’reilly, 2004; Mazzoni, et al., 2007), this may help explain why some prism adaptation studies have reported impaired error reduction while others have found reduced after-effects (Canavan, et al., 1990; Fernandez-Ruiz, et al., 2003; Stern, Mayeux, Hermann, & Rosen, 1988; Weiner, et al., 1983).

How is the prediction *p(outcome | movement plan)* generated, and how is the predictive model modified by the prism manipulation? The following two sections will offer a Bayesian account in answer to this question.

#### 4.3.1 Bayesian statistics and optimal state estimation

In order to generate appropriate motor commands given a movement plan, the nervous system needs to generate, update, and maintain accurate estimates of states of the body and the external world as we move through space (Franklin & Wolpert, 2011). During a pointing movement, multiple sources of error feedback (visual, proprioceptive, motor), with different time constants, both in-flight and following the reach endpoint, provide information about the hand and target position, and the discrepancy between the two. In addition, forward models generate advance predictions about expected outcomes before sensory observations occur. The nervous system must combine these multiple sources of information (predicted and observed) in order to generate a single integrated state estimate (e.g. position of the hand in space) that will guide behaviour (for review, see: Chandrasekaran, 2017; Ernst & Bülthoff, 2004). The uncertainty associated with any source of information places the problem of state estimation within a statistical framework. Bayesian statistics posit that inference about the state of the body or the external world can be made by combining and weighting each source of information according to its relative level of reliability. Consider for example the task of estimating the reach endpoint position of the hand after executing a pointing movement. Sensory predictions generated by forward models (e.g. proprioceptive forward model) constitute a prior assumption about where the hand should be if our internal models are correct. In Bayesian terms, such prior knowledge is represented as a *prior distribution* (blue curve in Figure 5A). The sensory system also provides (noisy) information about the likely position of the hand (e.g. proprioceptive feedback). This information is represented as a *likelihood distribution* (dotted black curve in Figure 5A). Combining both sources of information (prior and likelihood) according to Bayes’ rule offers a way to calculate a single, more certain estimate of the hand position, represented as a *posterior distribution* (red curve in Figure 5A). Because it minimises uncertainty, such estimate is called *optimal*.

**Figure 5.**
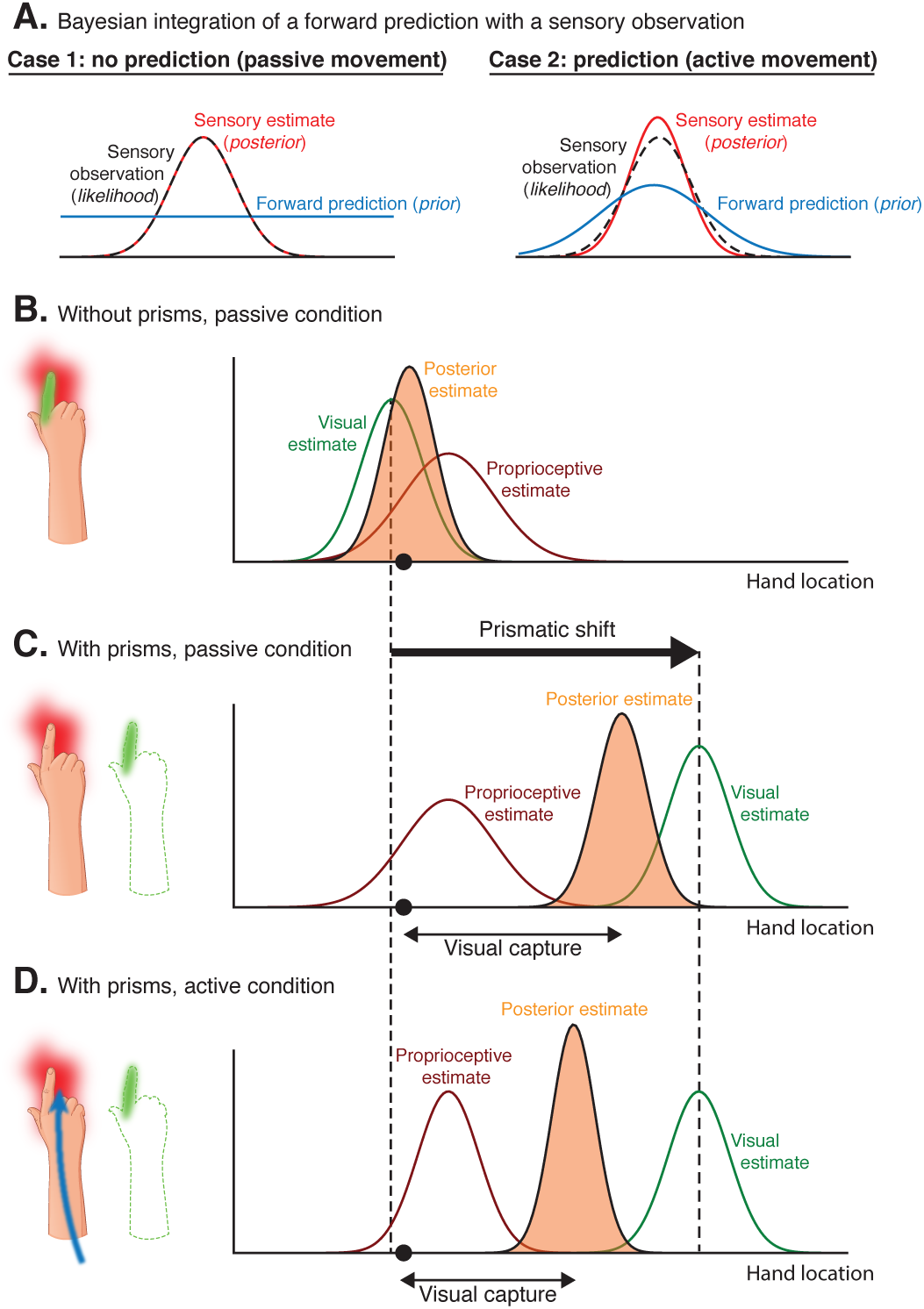
Bayesian statistics in sensorimotor control. Bayesian statistics describe how multiple sources of uncertain information can be combined optimally into a joint estimate. Here we consider the example of estimating the location of the hand in space. For all panels, the x-axis represents all possible locations of the hand and the y-axis is the probability of the hand being positioned at this location. **(A)** The optimal integration of a forward prediction (prior, in blue) with a sensory observation (likelihood, in black) is illustrated under two conditions. During a passive movement (case 1), no forward prediction is generated, i.e. the prior distribution is flat. The resulting sensory posterior estimate (in red) will therefore have the same distribution as the observation (likelihood). During an active movement however, a forward model generates a prediction of the next most likely hand position (prior) that can be integrated with sensory feedback (likelihood) to refine state estimation. This results in a posterior estimate (in red) that is more certain than the likelihood or the prior. **(B-D)** Bayesian statistics can also be used to describe (multimodal) visual-proprioceptive integration as illustrated here. The black dot on the x-axis represents the true location of the hand. **(B)** In the absence of prisms, the visual (green) and proprioceptive (red) estimates of the hand location are close to each other. Based on these two estimates, Bayes’ rule can be used to compute a posterior estimate of the hand location (purple) that is more certain than any of the sensory estimates. It posits that the relative level of uncertainty of the two sources of information (visual estimate and proprioceptive estimate) determines their relative contribution to the posterior estimate. Because the visual domain is more reliable (i.e. the width of the distribution is narrower), the resulting multimodal posterior estimate is biased towards the visual estimate. **(C)** The visuo-proprioceptive conflict induced by prism glasses is illustrated as a shift of the visual estimate (green) in the direction of the prismatic shift (towards the right). Bayes’ rule predicts that the resulting multimodal posterior estimate of the hand location is biased towards the visual observation because it is more reliable (i.e. lower standard deviation of the estimate) than the proprioceptive observation; this effect is called *visual capture*. The contribution of visual versus proprioceptive estimates of hand location to the multimodal posterior estimate is determined by their relative level of uncertainty. **(D)** In this condition, individuals actively move their limb without visual feedback prior to locating their hand in space. The execution of an active movement without visual feedback generates a proprioceptive forward prediction (case 2 of panel A) but no visual forward prediction (case 1 of panel A). If the proprioceptive forward prediction is accurate, the confidence in the resulting estimate will be increased. According to Bayes’ rule, the proprioceptive estimate will therefore have a greater influence on the multimodal posterior estimate of the hand location. In other words, the magnitude of visual capture is reduced.

Bayes’ rule states that the probability of the hand being in a location *x*, given a proprioceptive observation *o*, is the product of the prior probability of the hand location (i.e. prediction about the most likely position of the hand) and the likelihood (i.e. visual estimate of the hand location), normalized by the probability of the observation. Mathematically, Bayes’ rule defines the posterior as follow:

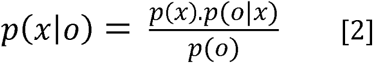

Several studies have shown that, when integrating multiple sources of uncertain information, humans behave in a nearly optimal way that is well predicted by Bayesian models (Alais & Burr, 2004; Ernst & Banks, 2002; Kayser & Shams, 2015; K. Kording & Wolpert, 2004; Konrad P Körding, et al., 2007; Sober & Sabes, 2005). Figure 5B illustrates the use of Bayes’ rule for visual-proprioceptive integration under the assumption of normally distributed noise. The uncertainty (i.e. width of the Gaussian curve) associated with the proprioceptive estimate has been found to be greater than that of its visual homologue (Beers, Sittig, & Denier, 1996; Burge, Girshick, & Banks, 2010; Ernst & Banks, 2002; van Beers, Sittig, & van Der Gon, 1999). Thus, according to Bayes’ rule, the optimal combination of proprioceptive and visual estimates of the hand location should result in a multimodal estimate that is biased towards the visual estimate.

In the context of prism adaptation, Bayesian statistics thus allows a re-formulation in mathematical terms of the perceptual phenomenon known as ‘visual capture’. Visual capture refers to the finding that individuals report their limb to be located closer to where it looks than where it feels under the visuo-proprioceptive conflict generated by prisms (Hay, Pick Jr, & Ikeda, 1965; Tastevin, 1937; van Beers, et al., 1999). Traditionally, this phenomenon has been described qualitatively as “a kind of perceptual fusion of the discrepant visual and proprioceptive stimuli”where “the proprioceptive stimulus […] fail[s] to evoke its normal response”(Hay, et al., 1965). By providing a rule to define optimal multisensory integration, Bayes’ theory makes quantitative predictions about the expected magnitude of sensory capture based on the level of uncertainty associated with information provided by each sensory modality (Burge, et al., 2010; Ernst & Banks, 2002; van Beers, et al., 1999) (Figure 5BC). It also offers a theoretical *explanation* for the reduction in magnitude of visual capture that has been observed following an active hand movement compared to a passive one (Welch, Widawski, Harrington, & Warren, 1979). The execution of a motor command elicits visual and proprioceptive forward models (efference copy) to predict the most likely next visual and proprioceptive observations (Lalazar & Vaadia, 2008). These predictions provide additional sources of information that can be combined with actual sensory observations to refine state estimation, i.e. reduce the uncertainty about the resulting percept. Because no visual feedback was provided during movement execution in Welch and colleagues’ experiment (Welch, et al., 1979), only the proprioceptive estimate of the hand position could benefit from integration of the prediction of a (proprioceptive) forward model. We suggest that this explains why the resulting percept was more biased towards the proprioceptive estimate (i.e. lower visual capture) in this condition (Figure 5D).

Visual capture has traditionally been studied relatively independently of prism adaptation, as it does not directly produce after-effects (Held & Hein, 1958; Welch, et al., 1979). The next section, however, will illustrate how, when expressed in Bayesian terms, the phenomenon of ‘visual capture’ is an important factor that will influence the relative extent of proprioceptive versus visual internal model updating that occurs during prism adaptation.

##### 4.3.2 Internal model updating and credit assignment

Prism after-effects occur only when active but not passive movements are executed under prism exposure (Held & Hein, 1958; Welch, et al., 1979). This is consistent with the idea that the generation of predictions prior to movement execution is fundamental to the ability to adapt in a way that generates after-effects. In a situation in which the reliability of sensory information is believed to be unchanged, persistent differences between predicted and measured observations implies the need for an update of internal models (Lalazar & Vaadia, 2008). The goal of this updating is to minimize systematic prediction errors and maintain internal consistency between predictions and observations.

In order to determine which internal model to update, the rate at which it should be updated, and for how long the update should be retained, the brain needs to solve what is known as a credit assignment problem, i.e. estimate the underlying *cause* of the prediction error, in order to assign learning to the correct internal model and update it appropriately (D. M. Wolpert, Diedrichsen, & Flanagan, 2011). It has been proposed that the way in which credit is assigned can be inferred behaviourally by predicted effects on: the rate of error reduction, or the pattern of generalization (to other contexts, spatial locations, body parts, modalities), or the timescale over which adaptation memory is retained (Max Berniker & Kording, 2008; Cothros, Wong, & Gribble, 2006; Kluzik, Diedrichsen, Shadmehr, & Bastian, 2008; K. P. Kording, et al., 2007; White & Diedrichsen, 2010).

When pointing towards visual targets while wearing prism glasses, there are many possible perturbations that could equally explain the occurrence of prediction errors. For example, a change in the arm dynamics – caused by fatigue for example – can alter the implementation of the motor command, causing systematic deviations from the predicted state. In this case, the nervous system should modify the relationship between a desired state and the associated motor command, i.e. adapt a *motor* internal model. Alternatively, prediction errors can arise from incorrect sensory estimates of the target location and/or of the position of the hand. In such a situation, the motor internal model does not necessarily need to be updated, but the *sensory* (visual and/or proprioceptive) internal models do.

Credit assignment refers to the complex problem of attributing a motor error to its causal source. It requires the nervous system to take into account a large number of parameters, which could include contextual cues, volatility of the environment, uncertainty in the estimates, predictions, and sensory observations, etc. Credit assignment may be solved by dedicated neural systems. For instance, computational models of visuomotor rotation that incorporate a Bayesian estimator of the source of the error signal have been able to successfully explain post-adaptation generalization (Max Berniker & Kording, 2008; Haith, Jackson, Miall, & Vijayakumar, 2009). Alternatively, in the absence of a dedicated estimator actively attempting to localize the error source, we suggest that a computational architecture that includes both modality-specific (visual and proprioceptive) and crossmodal levels of integration should be sufficient to allow the brain to sense which of the two internal models (visual or proprioceptive) is more likely to require an update. Figure 6 illustrates this architecture. For modality-specific levels of integration (i.e. visual prior/likelihood, proprioceptive prior/likelihood, see levels 7a and 7b of Figure 6), information theory offers a way to determine whether the prediction error (i.e. divergence between prior and likelihood) should be attributed to faulty forward or inverse models. Low confidence in the prediction (i.e. uncertain prior) and/or high confidence in the sensory observation (i.e. certain likelihood) imply the need for a forward model update. Alternatively, high confidence in the prediction (i.e. certain prior) and/or low confidence in the sensory observation (i.e. uncertain likelihood) are indicative of the need for an inverse model update. The comparison of the uncertainty in these visual versus proprioceptive estimates at the cross-modal level of integration (level 8 of Figure 6) should guide the neural system towards an update of the visual, proprioceptive or motor internal model. If both visual and proprioceptive estimates are similarly affected by the perturbation, it is likely to indicate an error in the inverse motor (i.e. the selected motor command was incorrect). However, in the case of a visuo-proprioceptive conflict (i.e. divergence between the visual and proprioceptive estimates), the sensory internal models associated with the most uncertain estimate (visual or proprioceptive) are the ones that should be preferentially updated. Because this computational architecture (Figure 6) incorporates the level of uncertainty associated with each source of information (width of the Gaussian curve) at every level of integration, there is sufficient information to determine which internal model (visual, proprioceptive and/or motor) should be updated, and how it should be updated.

Compared to the traditional dual-process theory, this framework makes predictions and offers a principled explanation of the predicted behavioural consequences of modulating the uncertainty of specific information sources (for example, see: Burge, et al., 2010; K. Kording & Wolpert, 2004; Yamamoto & Ando, 2012). For example, this can explain why terminal prism exposure (no visual feedback during reaching movements, endpoint error feedback only) induces greater proprioceptive than visual after-effects, while concurrent prism exposure (visual feedback during reach movements and endpoint error) induces greater visual than proprioceptive after-effects (Gordon M. Redding & Wallace, 1996; Gordon M Redding & Wallace, 2001, 2006b). The occlusion of visual feedback in terminal prism exposure conditions increases the uncertainty of the visual forward prediction of the endpoint position of the hand. Hence, when the prisms induce a prediction error, the cause of this is more likely to be attributed to a faulty visual internal model, thus resulting in greater visual than proprioceptive model updating. By contrast, the presence of visual feedback in concurrent prism exposure conditions allows for continuous state estimation to occur in both the visual and proprioceptive domains. It is known that in such condition that visual estimates are more certain relative to proprioceptive estimates (Burge, et al., 2010; Ernst & Banks, 2002, also see section 4.2.1 on visual capture and Figure 4). Hence, when a prediction error is experienced at the reach endpoint (under concurrent prism exposure), it is more likely to be attributed to a faulty proprioceptive internal model, which is then updated accordingly. Based on this interpretative framework, we predict that, across individuals, the magnitude of visual capture (which quantifies relative confidence in visual versus proprioceptive information) should predict the degree to which individuals update their visual relative to their proprioceptive internal models under concurrent prism exposure (i.e. the relative difference in after-effect magnitude between visual straight ahead judgement and proprioceptive straight ahead pointing).

**Figure 6.**
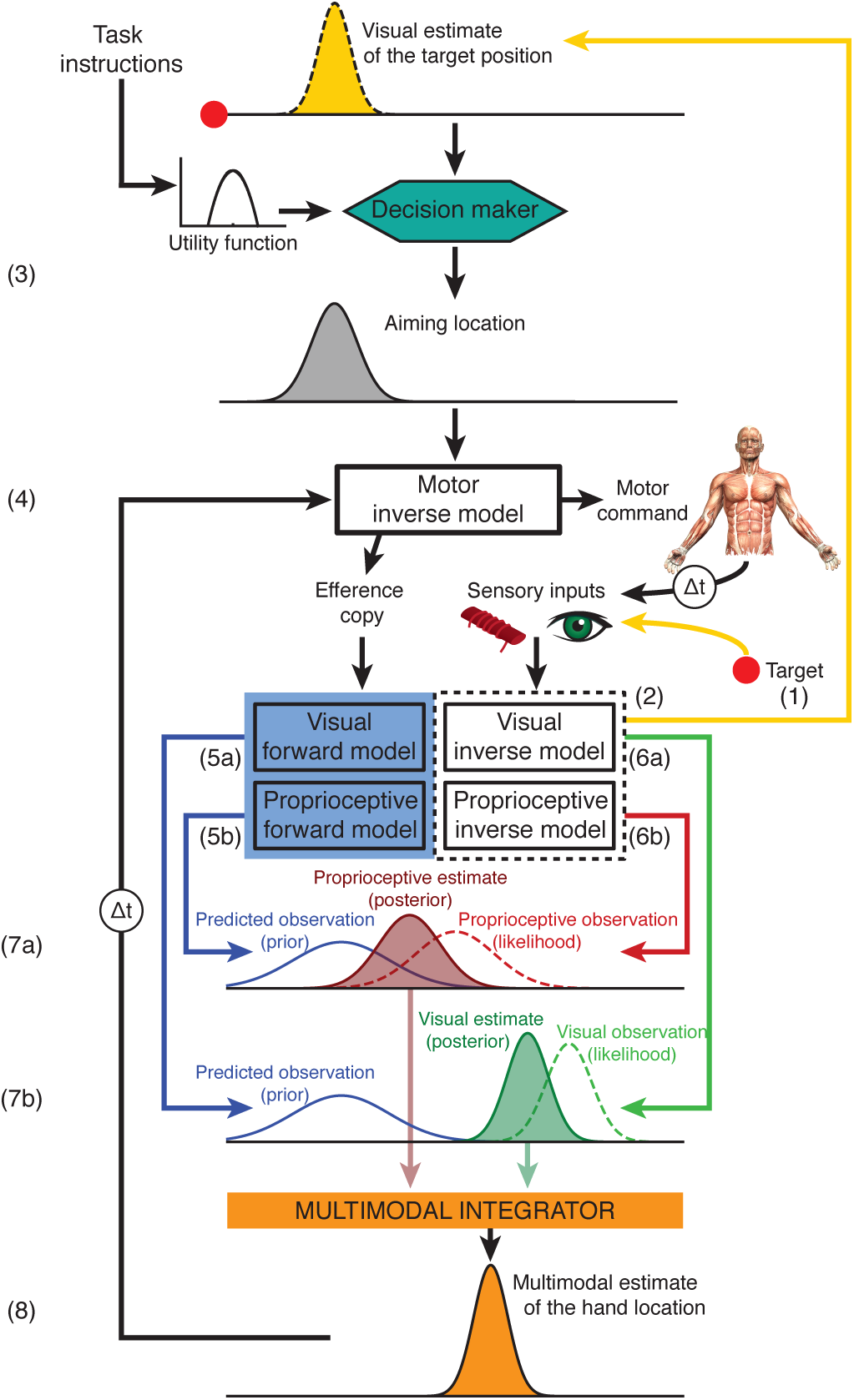
A Bayesian account of prism adaptation. This schema illustrates putative information processing occurring during the first reaching movement (closed loop pointing) on trial 1 of prism exposure. First, light from the target is refracted through the prism lens and enters the eye inducing sensory input (yellow arrow between the target and the eye, 1). Based on this input, a visual inverse model generates an estimate of the most likely location of the target (yellow arrow between the visual inverse model and the visual estimate of the target location, top of figure, 2). Owing to the prismatic displacement, this estimate will be right-shifted relative to the true location of the target in space (represented as the red dot, which is left of the visual estimate of the target location). The manual aiming direction believed to maximize expected utility, i.e. that which is expected to successfully align the reach endpoint with the target location, is selected by a decision maker (3). This process is explained in detail in Figure 4. The selected aiming location is then fed to a motor inverse model, which transforms the desired goal (aiming direction) into an action plan to accomplish it (motor command, 4). An efference copy of this motor command is sent to visual (5a) and proprioceptive (5b) forward models that generate modality-specific predictions about the most likely next location of the hand. In this example, because a ballistic movement is generated (closed loop pointing trial), this prediction concerns the hand position at the reach endpoint. Meanwhile, the execution of the motor command by the muscles generates visual and proprioceptive feedback (depicted by the arrows originating from the muscle spindle and eye symbols) that is integrated by sensory inverse models to generate modality-specific estimates of the sensed location of the hand (6a: visual and 6b: proprioceptive estimate of the hand location at the reach endpoint). Bayesian integration of the modality-specific forward predictions (prior) with the sensory observations (likelihood) generates modality-specific estimates of the most likely hand location (posterior) (7a and 7b). See Figure 5A for a detailed description of this type of integration. The resulting visual and proprioceptive (posterior) estimates are then combined into a multimodal (visuo-proprioceptive) posterior estimate of the hand position (8). See Figure 5B-D for a detailed description of this type of integration. On trial 1 of prism exposure, the rightward prism displacement induces: 1) leftward performance error (i.e. divergence between the visual estimate of the target position, 2, and the multimodal estimate of the hand location after movement execution, 8); 2) prediction errors in the sensory internal models (i.e. divergence between prior and likelihood, 7a-b); 3) visuo-proprioceptive conflict (i.e. divergence between the proprioceptive and visual estimates of the hand location, 8). The effect of internal model updating is to reduce these performance errors and to gradually realign all the distributions depicted in this schema (i.e. aiming location, predicted visual and proprioceptive observations, visual and proprioceptive observations, visual and proprioceptive estimates, multimodal estimate). Internal models are updated iteratively within this network as evidence accumulates and performance is adjusted from one trial to the next. In order to update internal models appropriately, the brain has to attribute learning to the correct internal models (motor, visual or proprioceptive). The level of uncertainty associated with each source of information (sensory predictions, sensory observations, sensory estimates) at every level of integration (modality-specific and multimodal) is used to infer the most likely error source and hence assign learning to the correct internal model. In this example, the visual estimate is more certain than the proprioceptive estimate, which should result in a preferential update of proprioceptive internal models. This model architecture incorporates the ‘strategic control versus spatial realignment’ dissociation as performance error can alternatively be quickly corrected at the level of the decision maker by selecting an aiming location that would result in a negative disparity between the visual estimate of the target position and the multimodal estimate of the reach endpoint hand location (see Figure 3 for a detailed description).

### 5 Conclusion

Prism adaptation is one of the oldest experimental paradigms used in the sensorimotor adaptation literature (Von Helmholtz, 1867). Yet, it has been studied mostly within a traditional ‘box-and-arrow’ cognitive psychology theoretical framework, which has important explanatory limitations. In this paper, we have advocated for the utility of a computational framework that re-conceptualizes prism adaptation in terms of its constituent algorithms. In our view, the advantages of a computational approach are several. First, Bayesian decision theory allows a precise quantitative re-formulation of ‘strategic control’ and ‘spatial realignment’ concepts, within the same utility framework used in other task contexts, such as reward-guided learning and decision-making (Behrens, et al., 2007; Daw, et al., 2006; Frank, et al., 2004). This shared conceptual and mathematical language opens up the possibility to investigate experimentally potential commonalities in the functional and neural mechanisms engaged across these very different task contexts. For example, one could hypothesize that there are some shared neural substrates responsible for aspects of the computation of expected utility notwithstanding the different types of action outcome experienced in these different classes of task. Second, Bayesian statistics provides a mathematical framework that specifies how spatial realignment should proceed, and thus offers quantitative explanation crucially lacking in the traditional framework. Third, state-space models offer a simple mathematical description of the temporal dynamics of internal models thought to drive prism adaptation behaviour, moving beyond the categorical dualist approach of the traditional framework. It enables quantitative questions about information processing and neural implementation, such as: 1) how many processes best explain prism adaptation behaviour?, and 2) is there a discrete number or a continuum of processes/timescales which varies with task parameters? 3) are distinct brain regions associated with distinct timescales, or might a given brain circuit have the capability to perform similar computations over a range of differing timescales?

To conclude, we submit that progress in understanding the functional and neural bases of prism adaptation behaviour requires this experimental paradigm to be reconceptualised at an algorithmic level of description. Doing so offers our field the opportunity to capitalize on explanatory gains generated by the literature on computational motor control. Such insights are being leveraged in the literature on other kinds of adaptation task, but the prism literature has so far remained oddly immune. Given the distinctive features of prism adaptation, and its applications in neuropsychology, we believe the time for our field to start leveraging these gains is ripe.

## Acknowledgements

PP was funded by the EU Marie Curie Initial Training Network (Adaptive Brain Computations). JO’R was funded by a Medical Research Council Career Development Award (MR/L019639/1). JO’S was funded by the Oxford Biomedical Research Centre.

## References

Aimola, L., Rogers, G., Kerkhoff, G., Smith, D. T., & Schenk, T. (2012). Visuomotor adaptation is impaired in patients with unilateral neglect. Neuropsychologia, 50, 1158–1163.

Alais, D., & Burr, D. (2004). The ventriloquist effect results from near-optimal bimodal integration. Current Biology, 14, 257–262.

Baizer, J. S., Kralj-Hans, I., & Glickstein, M. (1999). Cerebellar lesions and prism adaptation in macaque monkeys. J Neurophysiol, 81, 1960–1965.

Bedford, F. L. (1989). Constraints on learning new mappings between perceptual dimensions. Journal of Experimental Psychology: Human Perception and Performance, 15, 232.

Bedford, F. L. (1993). Perceptual and cognitive spatial learning. Journal of Experimental Psychology: Human Perception and Performance, 19, 517.

Beers, R. J., Sittig, A. C., & Denier, J. J. (1996). How humans combine simultaneous proprioceptive and visual position information. Experimental Brain Research, 111, 253–261.

Behrens, T. E., Woolrich, M. W., Walton, M. E., & Rushworth, M. F. (2007). Learning the value of information in an uncertain world. Nat Neurosci, 10, 1214–1221.

Bernacchia, A., Seo, H., Lee, D., & Wang, X.-J. (2011). A reservoir of time constants for memory traces in cortical neurons. Nat Neurosci, 14, 366–372.

Berniker, M., & Kording, K. (2008). Estimating the sources of motor errors for adaptation and generalization. Nat Neurosci, 11, 1454–1461.

Berniker, M., & Kording, K. (2011). Bayesian approaches to sensory integration for motor control. Wiley Interdiscip Rev Cogn Sci, 2, 419–428.

Bossom, J. (1965). The effect of brain lesions on prism-adaptation in monkey. Psychonomic Science, 2, 45–46.

Bultitude, J. H., Rafal, R. D., & Tinker, C. (2012). Moving forward with prisms: sensory-motor adaptation improves gait initiation in Parkinson’s disease. Frontiers in neurology, 3.

Burge, J., Girshick, A. R., & Banks, M. S. (2010). Visual– haptic adaptation is determined by relative reliability. Journal of Neuroscience, 30, 7714–7721.

Calzolari, E., Bolognini, N., Casati, C., Marzoli, S. B., & Vallar, G. (2015). Restoring abnormal aftereffects of prismatic adaptation through neuromodulation. Neuropsychologia, 74, 162–169.

Canavan, A. G. M., Passingham, R. E., Marsden, C. D., Quinn, N., Wyke, M., & Polkey, C. E. (1990). Prism adaptation and other tasks involving spatial abilities in patients with Parkinson’s disease, patients with frontal lobe lesions and patients with unilateral temporal lobectomies. Neuropsychologia, 28, 969–984.

Chandrasekaran, C. (2017). Computational principles and models of multisensory integration. Current Opinion in Neurobiology, 43, 25–34.

Chapman, H. L., Eramudugolla, R., Gavrilescu, M., Strudwick, M. W., Loftus, A., Cunnington, R., & Mattingley, J. B. (2010). Neural mechanisms underlying spatial realignment during adaptation to optical wedge prisms. Neuropsychologia, 48, 2595–2601.

Chaudhuri, R., Knoblauch, K., Gariel, M.-A., Kennedy, H., & Wang, X.-J. (2015). A large-scale circuit mechanism for hierarchical dynamical processing in the primate cortex. Neuron, 88, 419–431.

Clower, D. M., Hoffman, J. H., Votaw, J. R., Faber, T. L., Woods, R. P., & Alexander, G. E. (1996). Role of the posterior parietal cortex in the recalibration of visually guided reaching. Nature, 383, 618–621.

Colent, C., Pisella, L., Bernieri, C., Rode, G., & Rossetti, Y. (2000). Cognitive bias induced by visuo-motor adaptation to prisms: a simulation of unilateral neglect in normal individuals? Neuroreport, 11, 1899–1902.

Cothros, N., Wong, J., & Gribble, P. (2006). Are there distinct neural representations of object and limb dynamics? Experimental Brain Research, 173, 689–697.

Danckert, J., Ferber, S., & Goodale, M. A. (2008). Direct effects of prismatic lenses on visuomotor control: an event related functional MRI study. European Journal of Neuroscience, 28, 1696–1704.

Davidson, P. R., & Wolpert, D. M. (2005). Widespread access to predictive models in the motor system: a short review. Journal of Neural Engineering, 2, S313.

Daw, N. D., O’doherty, J. P., Dayan, P., Seymour, B., & Dolan, R. J. (2006). Cortical substrates for exploratory decisions in humans. Nature, 441, 876–879.

Dewar, R. (1971). Adaptation to displaced vision: Variations on the “prismatic-shaping” technique. Attention, Perception, & Psychophysics, 9, 155–157.

Ernst, M. O., & Banks, M. S. (2002). Humans integrate visual and haptic information in a statistically optimal fashion. Nature, 415, 429–433.

Ernst, M. O., & Bülthoff, H. H. (2004). Merging the senses into a robust percept. Trends in cognitive sciences, 8, 162–169.

Ethier, V., Zee, D. S., & Shadmehr, R. (2008). Spontaneous recovery of motor memory during saccade adaptation. J Neurophysiol, 99, 2577–2583.

Fernandez-Ruiz, J., Velasquez-Perez, L., Diaz, R., Drucker-Colin, R., Perez-Gonzalez, R., Canales, N., Sanchez-Cruz, G., Martinez-Gongora, E., Medrano, Y., Almaguer-Mederos, L., Seifried, C., & Auburger, G. (2007). Prism adaptation in spinocerebellar ataxia type 2. Neuropsychologia, 45, 2692–2698.

Fernandez-Ruiz, J., Diaz, R., Hall-Haro, C., Vergara, P., Mischner, J., Nunez, L., Drucker-Colin, R., Ochoa, A., & Alonso, M. (2003). Normal prism adaptation but reduced after-effect in basal ganglia disorders using a throwing task. European Journal of Neuroscience, 18, 689–694.

Frank, M. J., Seeberger, L. C., & O’reilly, R. C. (2004). By carrot or by stick: cognitive reinforcement learning in parkinsonism. Science, 306, 1940–1943.

Franklin, D. W., & Wolpert, D. M. (2011). Computational mechanisms of sensorimotor control. Neuron, 72, 425–442.

Frassinetti, F., Angeli, V., Meneghello, F., Avanzi, S., & Làdavas, E. (2002). Long-lasting amelioration of visuospatial neglect by prism adaptation. Brain, 125, 608–623.

Fusi, S., Drew, P. J., & Abbott, L. F. (2005). Cascade models of synaptically stored memories. Neuron, 45, 599–611.

Galea, J. M., Mallia, E., Rothwell, J., & Diedrichsen, J. (2015). The dissociable effects of punishment and reward on motor learning. Nat Neurosci, 18, 597–602.

Goedert, K. M., LeBlanc, A., Tsai, S.-W., & Barrett, A. M. (2010). Asymmetrical effects of adaptation to left-and right-shifting prisms depends on pre-existing attentional biases. Journal of the International Neuropsychological Society, 16, 795–804.

Haith, A., Jackson, C. P., Miall, R. C., & Vijayakumar, S. (2009). Unifying the sensory and motor components of sensorimotor adaptation. In Advances in Neural Information Processing Systems (pp. 593–600).

Hamilton, C. R. (1964). Intermanual transfer of adaptation to prisms. The American journal of psychology, 77, 457–462.

Hampton, A. N., Bossaerts, P., & O’doherty, J. P. (2006). The role of the ventromedial prefrontal cortex in abstract state-based inference during decision making in humans. Journal of Neuroscience, 26, 8360–8367.

Hanajima, R., Shadmehr, R., Ohminami, S., Tsutsumi, R., Shirota, Y., Shimizu, T., Tanaka, N., Terao, Y., Tsuji, S., Ugawa, Y., Uchimura, M., Inoue, M., & Kitazawa, S. (2015). Modulation of error-sensitivity during a prism adaptation task in people with cerebellar degeneration. J Neurophysiol, 114, 2460–2471.

Harris, C. M., & Wolpert, D. M. (1998). Signal-dependent noise determines motor planning. Nature, 394, 780–784.

Harris, C. S. (1963). Adaptation to displaced vision: visual, motor, or proprioceptive change? Science, 140, 812–813.

Hatada, Y., Miall, R., & Rossetti, Y. (2006). Two waves of a long-lasting aftereffect of prism adaptation measured over 7 days. Experimental Brain Research, 169, 417–426.

Hatada, Y., Miall, R. C., & Rossetti, Y. (2006). Long lasting aftereffect of a single prism adaptation: directionally biased shift in proprioception and late onset shift of internal egocentric reference frame. Experimental Brain Research, 174, 189–198.

Hatada, Y., Miall, R. C., & Rossetti, Y. (2006). Two waves of a long-lasting aftereffect of prism adaptation measured over 7 days. Experimental Brain Research, 169, 417–426.

Hatada, Y., Rossetti, Y., & Miall, R. C. (2006). Long-lasting aftereffect of a single prism adaptation: shifts in vision and proprioception are independent. Experimental Brain Research, 173, 415–424.

Hay, J. C., & Pick Jr, H. L. (1966). Visual and proprioceptive adaptation to optical displacement of the visual stimulus. Journal of experimental psychology, 71, 150.

Hay, J. C., Pick Jr, H. L., & Ikeda, K. (1965). Visual capture produced by prism spectacles. Psychonomic Science, 2, 215–216.

Held, R., & Hein, A. V. (1958). Adaptation of disarranged hand-eye coordination contingent upon re-afferent stimulation. Perceptual and Motor Skills, 8, 87–90.

Howard, I., & Freedman, S. (1968). Displacing the optical array. The neuropsychology of spatially oriented behavior, 19–36.

Hsu, M., Bhatt, M., Adolphs, R., Tranel, D., & Camerer, C. F. (2005). Neural systems responding to degrees of uncertainty in human decision-making. Science, 310, 1680–1683.

Huberdeau, D. M., Krakauer, J. W., & Haith, A. M. (2015). Dual-process decomposition in human sensorimotor adaptation. Current Opinion in Neurobiology, 33, 71–77.

Inoue, M., Uchimura, M., Karibe, A., O’Shea, J., Rossetti, Y., & Kitazawa, S. (2015). Three timescales in prism adaptation. J Neurophysiol, 113, 328–338.

Jacquin-Courtois, S., O’Shea, J., Luaute, J., Pisella, L., Revol, P., Mizuno, K., Rode, G., & Rossetti, Y. (2013). Rehabilitation of spatial neglect by prism adaptation: a peculiar expansion of sensorimotor after-effects to spatial cognition. Neurosci Biobehav Rev, 37, 594–609.

Joiner, W. M., & Smith, M. A. (2008). Long-term retention explained by a model of short-term learning in the adaptive control of reaching. J Neurophysiol, 100, 2948–2955.

Kawato, M. (1999). Internal models for motor control and trajectory planning. Current Opinion in Neurobiology, 9, 718–727.

Kayser, C., & Shams, L. (2015). Multisensory causal inference in the brain. PLoS Biol, 13, e1002075.

Kim, S., Ogawa, K., Lv, J., Schweighofer, N., & Imamizu, H. (2015). Neural Substrates Related to Motor Memory with Multiple Timescales in Sensorimotor Adaptation. PLoS Biol, 13, e1002312.

Kitazawa, S., Kimura, T., & Uka, T. (1997). Prism adaptation of reaching movements: specificity for the velocity of reaching. Journal of Neuroscience, 17, 1481–1492.

Kluzik, J., Diedrichsen, J., Shadmehr, R., & Bastian, A. J. (2008). Reach adaptation: what determines whether we learn an internal model of the tool or adapt the model of our arm? J Neurophysiol, 100, 1455–1464.

Kording, K., & Wolpert, D. M. (2004). Bayesian integration in sensorimotor learning. Nature, 427, 244–247.

Körding, K. P., Beierholm, U., Ma, W. J., Quartz, S., Tenenbaum, J. B., & Shams, L. (2007). Causal inference in multisensory perception. PLoS One, 2, e943.

Kording, K. P., Tenenbaum, J. B., & Shadmehr, R. (2007). The dynamics of memory as a consequence of optimal adaptation to a changing body. Nat Neurosci, 10, 779–786.

Körding, K. P., & Wolpert, D. M. (2004). The loss function of sensorimotor learning. Proc Natl Acad Sci U S A, 101, 9839–9842.

Körding, K. P., & Wolpert, D. M. (2006). Bayesian decision theory in sensorimotor control. Trends in cognitive sciences, 10, 319–326.

Kornheiser, A. S. (1976). Adaptation to laterally displaced vision: A review. Psychological bulletin, 83, 783.

Krakauer, J. W., Ghazanfar, A. A., Gomez-Marin, A., MacIver, M. A., & Poeppel, D. (2017). Neuroscience needs behavior: correcting a reductionist Bias. Neuron, 93, 480–490.

Krakauer, J. W., Pine, Z. M., Ghilardi, M.-F., & Ghez, C. (2000). Learning of Visuomotor Transformations for Vectorial Planning of Reaching Trajectories. The Journal of Neuroscience, 20, 8916–8924.

Kuper, M., Wunnemann, M. J., Thurling, M., Stefanescu, R. M., Maderwald, S., Elles, H. G., Goricke, S., Ladd, M. E., & Timmann, D. (2014). Activation of the cerebellar cortex and the dentate nucleus in a prism adaptation fMRI study. Hum Brain Mapp, 35, 1574–1586.

Lackner, J. R., & Dizio, P. (1994). Rapid adaptation to Coriolis force perturbations of arm trajectory. J Neurophysiol, 72, 299–313.

Lalazar, H., & Vaadia, E. (2008). Neural basis of sensorimotor learning: modifying internal models. Curr Opin Neurobiol, 18, 573–581.

Lee, J. Y., & Schweighofer, N. (2009). Dual adaptation supports a parallel architecture of motor memory. J Neurosci, 29, 10396–10404.

Loftus, A. M., Vijayakumar, N., & Nicholls, M. E. (2009). Prism adaptation overcomes pseudoneglect for the greyscales task. Cortex, 45, 537–543.

Luauté, J., Schwartz, S., Rossetti, Y., Spiridon, M., Rode, G., Boisson, D., & Vuilleumier, P. (2009). Dynamic Changes in Brain Activity during Prism Adaptation. The Journal of Neuroscience, 29, 169–178.

Martin, T., Keating, J., Goodkin, H., Bastian, A., & Thach, W. (1996). Throwing while looking through prisms. Brain, 119, 1183–1198.

Martin-Arevalo, E., Chica, A. B., & Lupianez, J. (2014). Electrophysiological modulations of exogenous attention by intervening events. Brain Cogn, 85, 239–250.

Mazzoni, P., Hristova, A., & Krakauer, J. W. (2007). Why don’t we move faster? Parkinson’s disease, movement vigor, and implicit motivation. Journal of Neuroscience, 27, 7105–7116.

Mazzoni, P., & Krakauer, J. W. (2006). An implicit plan overrides an explicit strategy during visuomotor adaptation. Journal of Neuroscience, 26, 3642–3645.

McDougle, S. D., Bond, K. M., & Taylor, J. A. (2015). Explicit and Implicit Processes Constitute the Fast and Slow Processes of Sensorimotor Learning. J Neurosci, 35, 9568–9579.

Miall, R. C., & Wolpert, D. M. (1996). Forward models for physiological motor control. Neural networks, 9, 1265–1279.

Michel, C., Pisella, L., Halligan, P. W., Luaute, J., Rode, G., Boisson, D., & Rossetti, Y. (2003). Simulating unilateral neglect in normals using prism adaptation: implications for theory. Neuropsychologia, 41, 25–39.

Michel, C., Pisella, L., Prablanc, C., Rode, G., & Rossetti, Y. (2007). Enhancing visuomotor adaptation by reducing error signals: single-step (aware) versus multiple-step (unaware) exposure to wedge prisms. Journal of Cognitive Neuroscience, 19, 341–350.

Morasso, P. (1981). Spatial control of arm movements. Experimental Brain Research, 42, 223–227.

Newport, R., & Jackson, S. R. (2006). Posterior parietal cortex and the dissociable components of prism adaptation. Neuropsychologia, 44.

O’Doherty, J. P., Dayan, P., Friston, K., Critchley, H., & Dolan, R. J. (2003). Temporal Difference Models and Reward-Related Learning in the Human Brain. Neuron, 38, 329–337.

O’doherty, J. P., Hampton, A., & Kim, H. (2007). Model-based fMRI and its application to reward learning and decision making. Ann N Y Acad Sci, 1104, 35–53.

O’Shea, J., Gaveau, V., Kandel, M., Koga, K., & Susami, K. (2014). Kinematic markers dissociate error correction from sensorimotor realignment during prism adaptation. Neuropsychologia.

Pisella, L., Michel, C., Grea, H., Tilikete, C., & Rossetti, Y. (2004). Preserved prism adaptation in bilateral optic ataxia: strategic versus adaptive reaction to prisms. Experimental Brain Research, 156.

Pisella, L., Rossetti, Y., Michel, C., Rode, G., Boisson, D., Pelisson, D., & Tilikete, C. (2005). Ipsidirectional impairment of prism adaptation after unilateral lesion of anterior cerebellum. Neurology, 65, 150–152.

Redding, G. M., Rader, S. D., & Lucas, D. R. (1992). Cognitive load and prism adaptation. Journal of motor behavior, 24, 238–246.

Redding, G. M., Rossetti, Y., & Wallace, B. (2005). Applications of prism adaptation: a tutorial in theory and method. Neuroscience & Biobehavioral Reviews, 29, 431–444.

Redding, G. M., & Wallace, B. (1988). Components of prism adaptation in terminal and concurrent exposure: Organization of the eye-hand coordination loop. Attention, Perception, & Psychophysics, 44, 59–68.

Redding, G. M., & Wallace, B. (1993). Adaptive Coordination and Alignment of Eye and Hand. Journal of motor behavior, 25, 75–88.

Redding, G. M., & Wallace, B. (1996). Adaptive spatial alignment and strategic perceptual-motor control. Journal of Experimental Psychology: Human Perception and Performance, 22, 379–394.

Redding, G. M., & Wallace, B. (2001). Calibration and alignment are separable: Evidence from prism adaptation. Journal of motor behavior, 33, 401–412.

Redding, G. M., & Wallace, B. (2002). Strategie Calibration and Spatial Alignment: A Model From Prism Adaptation. Journal of motor behavior.

Redding, G. M., & Wallace, B. (2006a). Generalization of prism adaptation. Journal of Experimental Psychology: Human Perception and Performance, 32, 1006.

Redding, G. M., & Wallace, B. (2006b). Prism adaptation and unilateral neglect: review and analysis. Neuropsychologia, 44, 1–20.

Rossetti, Y., Koga, K., & Mano, T. (1993). Prismatic displacement of vision induces transient changes in the timing of eye-hand coordination. Attention, Perception, & Psychophysics, 54, 355–364.

Rossetti, Y., Rode, G., Pisella, L., Farne, A., Boisson, D., & Perenin, M.-T. (1998). Prism adaptation to a rightward optical deviation rehabilitates left hemispatial neglect. Nature, 395, 166–169.

Schintu, S., Pisella, L., Jacobs, S., Salemme, R., Reilly, K. T., & Farne, A. (2014). Prism adaptation in the healthy brain: the shift in line bisection judgments is long lasting and fluctuates. Neuropsychologia, 53, 165–170.

Serino, A., Barbiani, M., Rinaldesi, M. L., & Làdavas, E. (2009). Effectiveness of prism adaptation in neglect rehabilitation. Stroke, 40, 1392–1398.

Shadmehr, R., Huang, H. J., & Ahmed, A. A. (2016). A representation of effort in decision-making and motor control. Current Biology, 26, 1929–1934.

Shadmehr, R., & Mussa-Ivaldi, F. A. (1994). Adaptive representation of dynamics during learning of a motor task. Journal of Neuroscience, 14, 3208–3224.

Shadmehr, R., Smith, M. A., & Krakauer, J. W. (2010). Error correction, sensory prediction, and adaptation in motor control. Annu Rev Neurosci, 33, 89–108.

Simmons, G., & Demiris, Y. (2005). Optimal robot arm control using the minimum variance model. Journal of Field Robotics, 22, 677–690.

Smith, M. A., Ghazizadeh, A., & Shadmehr, R. (2006). Interacting adaptive processes with different timescales underlie short-term motor learning. PLoS Biol, 4, e179.

Sober, S. J., & Sabes, P. N. (2005). Flexible strategies for sensory integration during motor planning. Nat Neurosci, 8, 490–497.

Stern, Y., Mayeux, R., Hermann, A., & Rosen, J. (1988). Prism adaptation in Parkinson’s disease. Journal of Neurology, Neurosurgery & Psychiatry, 51, 1584–1587.

Striemer, C. L., Russell, K., & Nath, P. (2016). Prism adaptation magnitude has differential influences on perceptual versus manual responses. Experimental Brain Research, 234, 2761–2772.

Sumitani, M., Rossetti, Y., Shibata, M., Matsuda, Y., Sakaue, G., Inoue, T., Mashimo, T., & Miyauchi, S. (2007). Prism adaptation to optical deviation alleviates pathologic pain. Neurology, 68, 128–133.

Sutton, R. S., & Barto, A. G. (1998). Reinforcement learning: An introduction (Vol. 1): MIT press Cambridge.

Tastevin, J. (1937). En partant de l’expérience d’Aristote les déplacements artificiels des parties du corps ne sont pas suivis par le sentiment de ces parties ni par les sensations qu’on peut y produire. L’Encéphale: Revue de psychiatrie clinique biologique et thérapeutique.

Templeton, W. B., Howard, I. P., & Wilkinson, D. A. (1974). Additivity of components of prismatic adaptation. Attention, Perception, & Psychophysics, 15, 249–257.

Trommershäuser, J., Maloney, L. T., & Landy, M. S. (2003). Statistical decision theory and the selection of rapid, goal-directed movements. JOSA A, 20, 1419–1433.

van Beers, R. J., Sittig, A. C., & van Der Gon, J. J. D. (1999). Integration of proprioceptive and visual position-information: An experimentally supported model. J Neurophysiol, 81, 1355–1364.

Von Helmholtz, H. (1867). Handbuch der physiologischen Optik (Vol. 9): Voss.

Weiner, M. J., Hallett, M., & Funkenstein, H. H. (1983). Adaptation to lateral displacement of vision in patients with lesions of the central nervous system. Neurology, 33, 766–766.

Welch, R. B., & Goldstein, G. (1972). Prism adaptation and brain damage. Neuropsychologia, 10, 387–394.

Welch, R. B., Widawski, M. H., Harrington, J., & Warren, D. H. (1979). An examination of the relationship between visual capture and prism adaptation. Attention, Perception, & Psychophysics, 25, 126–132.

White, O., & Diedrichsen, J. (2010). Responsibility assignment in redundant systems. Current Biology, 20, 1290–1295.

Wolpert, D. M., Diedrichsen, J., & Flanagan, J. R. (2011). Principles of sensorimotor learning. Nat Rev Neurosci, 12, 739–751.

Wolpert, D. M., & Kawato, M. (1998). Multiple paired forward and inverse models for motor control. Neural networks, 11, 1317–1329.

Yamamoto, M., & Ando, H. (2012). Probabilistic models of state estimation predict visuomotor transformations during prism adaptation. Vis Neurosci, 29, 119–129.

Zarahn, E., Weston, G. D., Liang, J., Mazzoni, P., & Krakauer, J. W. (2008). Explaining savings for visuomotor adaptation: linear time-invariant state-space models are not sufficient. J Neurophysiol, 100, 2537–2548.

